# *In Situ* Optical Quantification of Extracellular Electron Transfer using Plasmonic Metal Oxide Nanocrystals

**DOI:** 10.1101/2020.10.13.336008

**Authors:** Austin J. Graham, Stephen L. Gibbs, Camila A. Saez Cabezas, Yongdan Wang, Allison M. Green, Delia J. Milliron, Benjamin K. Keitz

## Abstract

Extracellular electron transfer (EET) is a critical form of microbial metabolism that enables respiration on a variety of inorganic substrates, including metal oxides. However, quantifying current generated by electroactive bacteria has been predominately limited to biofilms formed on electrodes. To address this, we developed a platform for quantifying EET flux from cell suspensions using aqueous dispersions of infrared plasmonic tin-doped indium oxide nanocrystals. Tracking the change in optical extinction during electron transfer enabled quantification of current generated by planktonic *Shewanella oneidensis* cultures. Using this method, we differentiated between starved and actively respiring cells, between cells of varying genotype, and between cells engineered to differentially express a key EET gene using an inducible genetic circuit. Overall, our results validate the utility of colloidally stable plasmonic metal oxide nanocrystals as quantitative biosensors in native biological environments and contribute to a fundamental understanding of planktonic *S. oneidensis* electrophysiology using simple *in situ* spectroscopy.

## Introduction

Extracellular electron transfer (EET) is a microbial respiratory process by which electron flux from carbon metabolism is transported across the cell membrane and directed onto inorganic substrates (1). This capability makes electrogenic bacteria useful organisms for microbial fuel cells (2,3), bioelectronics (4–6), material syntheses (7, 8), and has led to significant interest in controlling their electron flux using synthetic biology (9, 10). Improving EET-dependent technologies such as controlling electron transfer dynamics (11), implementing genetic logic (12), and orchestrating electron transfer dynamics between proteins (13) necessitates a stronger quantitative understanding of bacterial electrophysiology and simple methods to measure current generation. EET is typically quantified from biofilms formed on electrodes in microbial fuel cells (MFCs) or bioelectrochemical systems (BESs) (1, 14–16). EET from single cells has also been measured in such systems, providing sensitive quantitative measurements and highlighting variability in EET capabilities between individual cells (17–19). However, these measurements often require long incubation times and electrode colonization (∼days), and the contribution of suspended (*i*.*e*., planktonic) microbes is generally unclear. Furthermore, implementation of synthetic gene programs such as genetic logic operations often require cell turnover and growth, which are more complex in biofilms compared to planktonic culture. While there has been recent progress toward quantifying EET from planktonic cells in BESs (20), EET from planktonic microbes is more traditionally quantified optically, using soluble substrates such as dye-bound iron citrate or riboflavin (12, 21–23). We propose that by expanding upon the optical analysis toolkit by employing dispersed, plasmonic nanoparticles, we will enable greater understanding of EET to insoluble substrates.

Plasmonic nanoparticles, though insoluble, maintain colloidal stability in a variety of solvents. An intrinsically high concentration of conductive electrons results in a strong absorption at the localized surface plasmon resonance (LSPR) peak frequency, *ω*_*LSPR*_, and enables participation in redox reactions with molecules near the surface many times over. The LSPR peak position and intensity depend on the particle’s conduction electron concentration, *n*_*e*_, such that tracking the optical response near *ω*_*LSPR*_ can report on the number of redox reactions occurring at the nanoparticle surface (24–26). Thus, optically tracking the plasmonic response can monitor electron transfer similar to tracking current from a traditional film electrode. Doped metal oxide nanocrystals are particularly promising candidates because they sustain plasmonic resonance but have *n*_*e*_ ∼1-3 orders of magnitude lower than Au and Ag (27, 28), making their optical response more sensitive to single electron transfer events and also placing *ω*_*LSPR*_ in the infrared (24, 25).

Doped metal oxide nanocrystals have a few advantages over traditional optically-monitored acceptors such as iron citrate and flavins. First, *ω*_*LSPR*_ can be tuned throughout the near- to mid-infrared. This enables tailoring the peak absorption to the ideal window for each experimental setting, and leaves the visible spectrum available for separate and simultaneous measurements. Further, the surface of doped metal oxide nanocrystals can be post-synthetically functionalized with various ligands to customize solvent compatibility and tune the interactions with other species in solution (29–31). In addition, a large insoluble acceptor should inhibit passive and active transport across the cell membrane (32), both of which could convolute reduction pathways when using small molecule acceptors such as iron citrate or flavins. For example, microbial iron reduction does not necessarily translate to electricity generation on an electrode from that organism (33).

Overall, there is a need for reliable *in situ* biosensors of EET that are compatible with planktonic culture. Indeed, the advantageous properties of metal oxide nanocrystals have already enabled their use in spectroscopic measurement of EET (34–37). However, these methods did not directly quantify the relationship between the spectroscopic response and the electron transfer rate (*i*.*e*., current generation). We recently developed a model for fitting the plasmonic optical response of doped metal oxide nanocrystal dispersions which extracts the number of free electrons in each nanocrystal, enabling *in situ* tracking of biological electron transfer events using infrared spectroscopy (38).

Leveraging our ability to quantify the change in optical response with the aqueous biocompatibility of colloidal nanocrystal dispersions, we aimed to quantify *in situ* electron flux from planktonic microbes in culture suspensions. We chose *Shewanella oneidensis* as a model electroactive bacterium that is attractive for its respiratory plasticity and genetic tractability. Using non-destructive optical measurements to track the LSPR peak in real-time, we successfully quantified kinetic EET rate constants from *S. oneidensis* of varying metabolic activity and genotype. We also quantitatively differentiated electron flux from an engineered *S. oneidensis* strain with tuned EET gene expression levels. Our method enabled quantification of *S. oneidensis* population currents from planktonic culture, with the average cell exhibiting 0.05 – 2.8 fA • cell^−1^ depending on metabolic activity, genotype, and gene expression level. Overall, our results indicate that plasmonic semiconductor nanocrystals are a reliable infrared sensing platform for probing metabolic activity in cellular environments and contribute to a quantitative understanding of EET.

## Results & Discussion

### Preparation of Sn-doped indium oxide (ITO) nanocrystal aqueous dispersions

ITO nanocrystals (5.78 ± 0.64 nm radius, 9.4 at% Sn) were synthesized using established colloidal methods and a slow injection approach (Figure 1a, Materials and Methods) (39). Nanocrystal size and Sn doping concentration were measured with small-angle X-ray scattering (SAXS) (Figure S1) and inductively coupled plasma-optical emission spectroscopy, respectively. To disperse ITO nanocrystals in aqueous solvents, the hydrophobic organic ligands bound to the surface of the as-synthesized nanocrystals were replaced by a hydrophilic polymer. Specifically, we chose poly(acrylic acid) grafted with methoxy-terminated poly(ethylene oxide) (PAA-mPEO_4_). This custom-designed and biocompatible random co-polymer has previously been used to functionalize different types of metal oxides (40, 41) and been shown to be suitable for facilitating electrochemical redox reactions in thin films of polymer-wrapped ITO nanocrystals (42).

**Figure 1.**
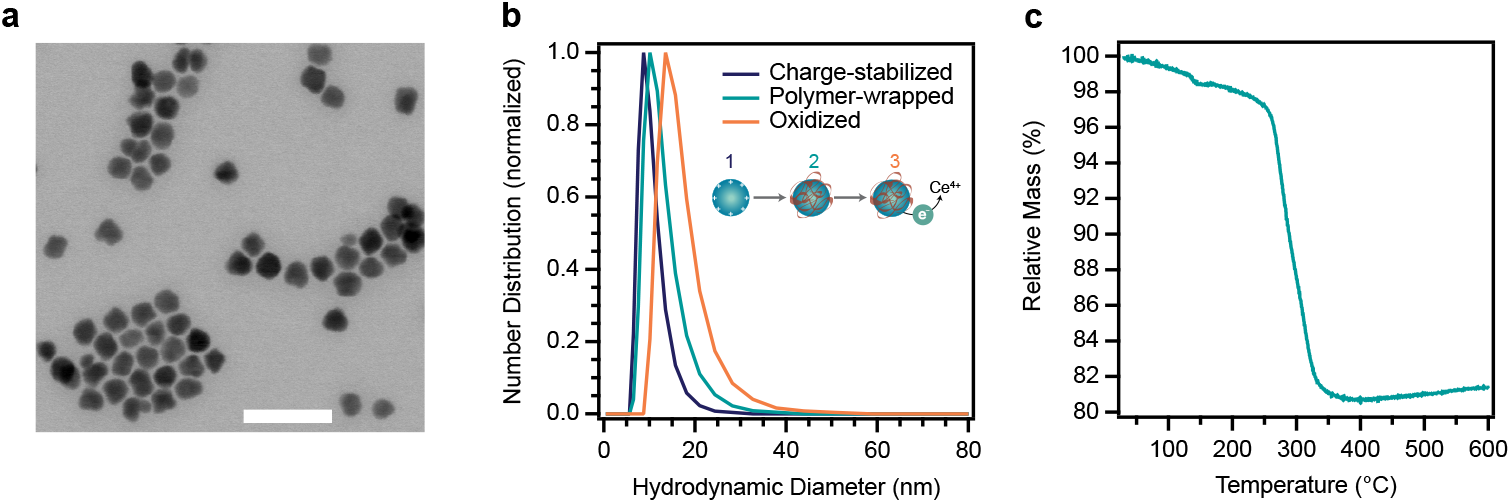
Preparation of polymer-wrapped oxidized nanocrystals for EET biosensing. (a) Scanning transmission electron microscopy image of as-synthesized ITO nanocrystals. Scale bar = 40 nm. (b) Dynamic light scattering of charge-stabilized, polymer-wrapped, and oxidized polymer-wrapped ITO nanocrystals. The ITO nanocrystals remain colloidally stable and do not exhibit signs of significant aggregation after each processing step (inset) since the hydrodynamic diameter measurements of the three samples are in close agreement. (c) Quantification of polymer adsorption on the ITO nanocrystal surface using thermogravimetric analysis. The mass loss (18%) around 300 °C corresponds to the decomposition of PAA-mPEO_4._

The functionalization of ITO nanocrystals with PAA-mPEO_4_ was achieved in a two-step process. First, the native stabilizing ligands were chemically removed by exposing the ITO nanocrystals to nitrosonium tetrafluoroborate, which resulted in bare, charge-stabilized ITO nanocrystals dispersible in N,N-dimethylformamide (31). Then, charge-stabilized ITO nanocrystals were mixed with PAA-mPEO_4_ and subsequently transferred into Milli-Q water. In a previous study, we found that colloidal stability in aqueous media is due to the adsorption of PAA-mPEO_4_ on ITO and is likely facilitated by the coordination of carboxylate species at the nanocrystal surface (42). Fourier transform infrared spectroscopy of nanocrystal dispersions at each processing step confirmed that the native ligands were effectively removed and replaced by PAA-mPEO_4_ (Figure S2). Finally, polymer-wrapped ITO nanocrystals were chemically oxidized upon the addition of ammonium cerium (IV) nitrate to promote EET from *S. oneidensis*. As expected, oxidation of the nanocrystal dispersion caused a red-shift and decrease in intensity of the LSPR peak (Figure S3a). After oxidation, the polymer-wrapped ITO nanocrystals were dispersed in a deuterated medium (*Shewanella* Basal Medium, SBM, Table S1). Deuterated buffer avoided saturating the spectrophotometer detector, as strong water absorption in the near-infrared (NIR) (43) overlaps significantly with the ITO LSPR peak (Note S1). A non-deuterated buffer could in principle be used by employing a doped metal oxide nanocrystal material with higher energy *ω*_*LSPR*_, such as F, In co-doped CdO (27).

Avoiding the formation of large ITO nanocrystal aggregates in the dispersion before or during EET is key for interpretation and analysis of optical measurements. Therefore, we monitored the hydrodynamic diameter of the nanocrystal dispersion after each processing step (*i*.*e*., ligand removal, polymer wrapping, and oxidation post-polymer wrapping) using dynamic light scattering to probe the stability of the colloid (Figure 1b). The hydrodynamic diameters of the charge-stabilized and polymer-wrapped samples were centered around ∼10 nm, which is in good agreement with the diameter obtained from SAXS. The slight increase in the hydrodynamic diameter of the polymer-wrapped and oxidized sample (D_h_ ∼15 nm) could suggest interaction between the colloid and charged species during oxidation, but we did not detect signs of significant aggregation. This dispersion was colloidally stable for at least five months after oxidation and transfer into deuterated SBM (Figure S3b). We hypothesize that the colloidal stability of our aqueous dispersions of polymer-wrapped ITO (PAAPEO-ITO) is due to sufficient PAA-mPEO_4_ adsorption on the nanocrystal surface (18% by mass, Figure 1c), which we quantified by thermogravimetric analysis, and thus effective steric stabilization.

### *S. oneidensis* remains viable in the presence of PAAPEO-ITO

All bacterial strains and plasmids used in this study are listed in Table S3. We first ensured that polymer-wrapped nanocrystals were not cytotoxic by quantifying cell viability in their presence (44). Aerobically pregrown cells were diluted to OD_600_ = 0.2 in a mixture of ∼1 mg/mL (∼0.2 µM) PAAPEO-ITO and aerobically incubated for 2 h at 30 °C. Cells were then stained for membrane permeability using the BacLight Live/Dead kit (Invitrogen) according to manufacturer protocols. Unbound Live/Dead stain was washed away and cells were imaged using green (live) and red (dead) fluorescence channels. Under these conditions, *S. oneidensis* MR-1 (wild-type) remained predominately alive, with an 86 ± 2% viable population (Figure S4). To assess cell health for longer experiments, colony counting was used to quantify the viable population after 8 h incubation with PAAPEO-ITO and 20 mM lactate at room temperature in deuterated SBM with casamino acids. As expected, there were not significant effects on the viable population (Figure S5). Minor decreases in viable population can likely be attributed to the lack of electron acceptor aside from PAAPEO-ITO, which was present at ∼1.5 mg/mL (∼0.3 µM) for EET quantification experiments (compared to standard growth conditions in 40 mM fumarate). In contrast to fumarate, a nanocrystal can accept more than one electron, but not enough to account for this concentration difference. These results indicate that cell viability is not substantially impacted beyond electron acceptor limitation, and that polymer-wrapped nanocrystals are not cytotoxic to *S. oneidensis*.

### Optical extinction of PAAPEO-ITO tracks respiratory electron flux from *S. oneidensis*

Once cell viability was established, we measured the *in situ* optical response of the PAAPEO-ITO LSPR during electron transfer from *S. oneidensis* MR-1. Aerobically pregrown cells were washed and resuspended in 1.5 mg/mL PAAPEO-ITO in deuterated SBM with casamino acids and 20 mM lactate as the carbon source. The solution was immediately transferred to a cuvette, sealed, and loaded into a spectrophotometer in transmission mode to monitor the LSPR peak over time.The time-resolved increase in optical density and blue shift of the LSPR peak results from an increasing number of electrons within the ITO NCs, which indicates continuous electron transfer to the nanocrystals (Figure 2a). These results confirm that *S. oneidensis* is able to reduce PAAPEO-ITO and the kinetics of this process are resolvable using NIR spectroscopy. Given the aerobic preparation of the cell culture and sample, they also suggest that dissolved oxygen is first consumed through aerobic respiration, followed by activation of anaerobic EET pathways (*i*.*e*., reduction of PAAPEO-ITO), as we have previously observed for other electron acceptors such as copper and fumarate (45, 46). We then measured EET from a *S. oneidensis* MR-1 population deprived of a carbon source (*e*.*g*., lactate), which should inhibit respiration and prevent generation of electron flux. The change in optical extinction was significantly diminished in these samples (Figure 2b). Some minimal reduction was observed, which can likely be attributed to excess metabolism from pregrowth (7), or to non-specific reducing capacity from cell stress and lysis. Indeed, a reaction containing lysed *S. oneidensis* slightly reduced PAAPEO-ITO even without oxygen removal, indicating that cell death could lead to a small amount of nonspecific reduction (Figure S6). Similarly, an *E. coli* sample demonstrated minimal PAAPEO-ITO reduction, as would be expected from this EET-deficient microbe (Figure 3c). Together, these measurements confirm that regulating metabolic flux manifests in an observable difference to the optical response of PAAPEO-ITO, validating this platform as an effective *in situ* biosensor that is specific to EET.

**Figure 2.**
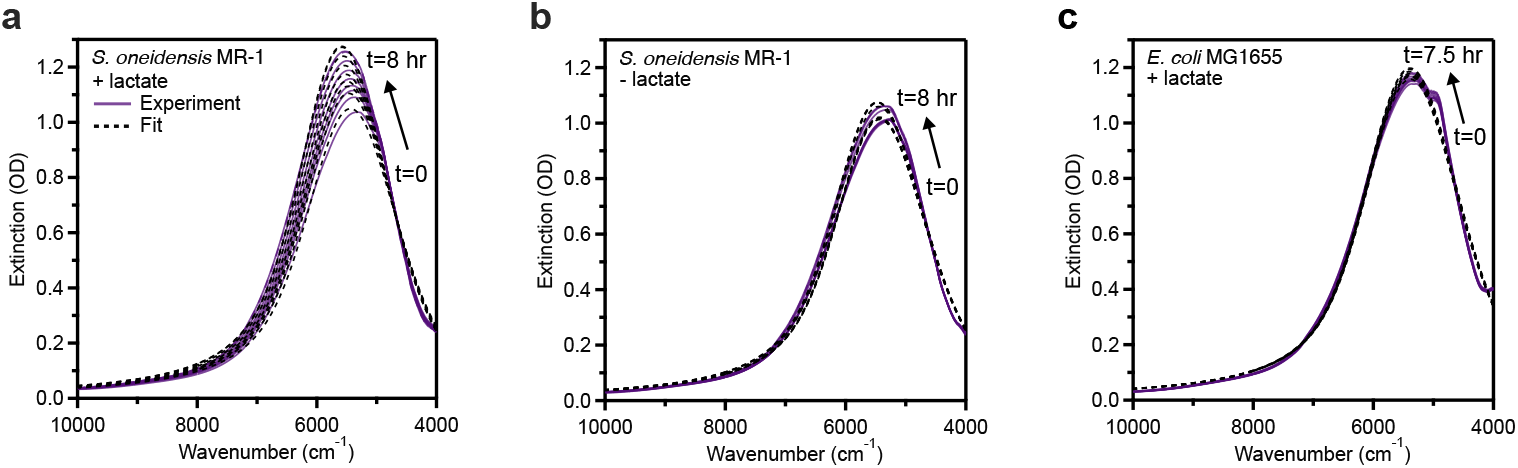
EET from metabolically active *S. oneidensis* leads to time-resolved increased optical extinction of PAAPEO-ITO. Experimental extinction spectra collected each hour from *t* = 0 to 8 h and associated fits for PAAPEO-ITO inoculated with (a) *S. oneidensis* MR-1 that was provided 20 mM lactate as a carbon source, (b) *S. oneidensis* MR-1 not provided a carbon source, or (c) *E. coli* MG1655 provided with 20 mM lactate as a carbon source.

**Figure 3.**
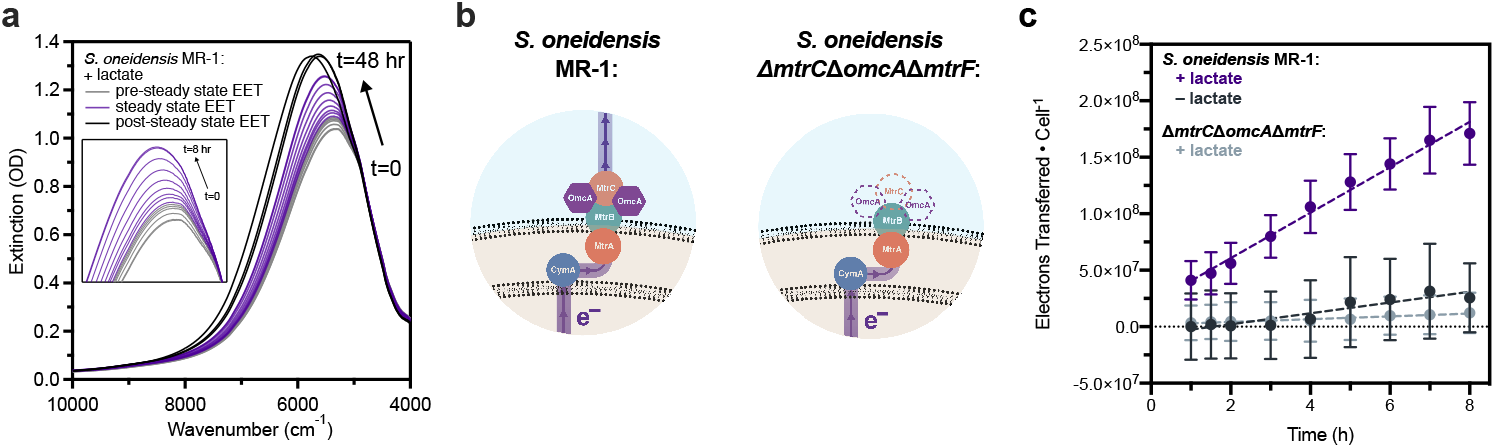
HEDA fitting of PAAPEO-ITO LSPR response during EET from *S. oneidensis* enables quantification of steady-state electron transfer kinetics. (a) EET from *S. oneidensis* to PAAPEO-ITO exhibits three domains. (Inset) The early domain (≤ 1 h) shows variability due to aerobic respiration, introduction of a deuterated solvent, and cell stress. This regime includes a lag phase (0–20 min) followed by a sharp increase in electron transfer (20–60 min). Steady-state electron transfer (1–8 h) precedes a later domain (≥ 8 h) exhibiting effects of nanocrystal saturation, precipitation, and cell death. (b) The Mtr pathway in *S. oneidensis*. In the Δ*mtrC*Δ*omcA*Δ*mtrF* knockout (right), three critical extracellular cytochrome genes are removed from the genome, hindering EET capabilities. (c) Steady-state EET (1–8 h) where dashed lines represent linear regression of the electrons transferred over time and *k*_*EET*_ is the slope of the regression (Table 1). Data points shown are the mean ± SD of *n* = 3 fits of biological replicates.

As a proof of concept, we also collected extinction out to higher energies to test if we could simultaneously measure the infrared LSPR spectrum and meaningful data in the visible wavelengths. We chose to quantify the intensity of scattered light by the cells at 600 nm (OD_600_) because this value is used to quantify the concentration of cells in a dispersion. We were able to successfully monitor both the infrared plasmon peak and the OD_600_ value in a single sweep of wavelengths in our spectrophotometer (Figure S7), further illustrating the benefits of developing an infrared sensing platform.

### Quantitative spectral analysis enables quantification of EET from *S. oneidensis* with varying metabolic activity and genotype

Following qualitative observations of the LSPR peak over time, we fit the peaks to a free electron model that extracts the number of conductive electrons in a nanoparticle. Using a deuterated buffer allowed for the collection of clean spectra and nearly the full plasmon peak for the ITO nanocrystals could be resolved. ITO nanocrystals were previously incubated with *S. oneidensis*, but only to quantify total electrons transferred from wild-type *S. oneidensis*, and only in non-deuterated buffer (36). Therein the water absorption obscured the signal below ∼5500 cm^−1^, meaning the high energy tail of the LSPR peak and the ultraviolet band gap absorption were used to quantify electron transfer. While that work was an important step towards quantifying EET using ITO nanocrystals, our ability to resolve nearly the full plasmonic response allows for a simpler, more reliable quantification using our recently developed model, the heterogeneous ensemble Drude approximation (HEDA). The HEDA model extracts intrinsic material properties, such as electron concentration, from the plasmonic response of a dilute dispersion of nanocrystals (38). By accounting for ensemble heterogeneities in nanocrystal size and nanocrystal electron concentration, it enables more accurate acquisition of material properties.

Immediately upon inoculating the cells with oxidized PAAPEO-ITO, we began collecting time-resolved spectra, continuing for up to 48 hours. We fit spectra using the HEDA model to quantify the number of electrons within the average nanocrystal in an ensemble at each timepoint (Note S2). Since we quantified both the concentration of nanocrystals and cells, we could calculate the cumulative electrons transferred per cell over time. This quantification revealed two consistent electron transfer regimes across samples – a variable regime during the first ∼1 h, where electron transfer was nonlinear (Figure 3a, inset; Figure S8a), followed by a steady-state electron transfer regime. In previous studies, aerobic *S. oneidensis* at OD_600_ = 0.2 took between 10 min to 1 h to fully consume dissolved oxygen and turn on anaerobic EET pathways, depending on the electron acceptor and reaction conditions (45, 46). Therefore, we attribute this initial variable domain to a combination of oxygen removal via aerobic respiration and cell stress as they adjust to a new electron acceptor and undergo a metabolic shift.

Beyond 8 h of reaction in high EET samples (*e*.*g*., wild-type *S. oneidensis* with lactate), we noticed effects of nanocrystal saturation and precipitation (Figure 3a, Figure S8b). Thus, we defined the steady-state regime as between 1 and 8 h, and used this domain to determine electron transfer kinetics. Notably, OD_600_ = 0.2 is approximately the saturation density of anaerobic *S. oneidensis* in SBM with 20 mM lactate, although this varies depending on the electron acceptor. We suspect that upon adjusting to their new anaerobic environment, the cells should divert minimal metabolic flux toward growth and instead direct it toward energy production via EET (47). This, combined with the lack of cell growth between inoculation and 8 h (Figure S5), validates the appearance of steady-state electron transfer in our system. As such, the linear increase in electrons transferred per cell justified using linear regressions to obtain kinetic rate constants and compute population-averaged current generation. EET could also be measured in samples containing an added electron acceptor, fumarate. However, to simplify analysis and ensure all metabolic electron flux was directed toward the measured acceptor, PAAPEO-ITO, we chose to omit fumarate from future experiments (Figure S9).

We first determined steady-state electron transfer rates to PAAPEO-ITO for *S. oneidensis* MR-1 with and without lactate as a carbon source, which qualitatively showed a large difference in optical response (Figure 2). When 20 mM lactate was present as a carbon source, enabling continuous respiration and generation of EET flux, the rate of electron transfer was *k*_*EET*_ = 2.0 ± 0.2 × 10^7^ electrons • cell^−1^ • h^−1^, or 0.89 ± 0.07 fA • cell^−1^, which is approximately 5-fold greater than when the cells were starved (Table 1). To further assess background electron transfer and dynamic range, we used an EET-deficient strain of *S. oneidensis. S. oneidensis* Δ*mtrC*Δ*omcA*Δ*mtrF* is a triple-knockout of three key extracellular cytochromes responsible for direct electron transfer to a variety of substrates (Figure 3b), and as a result, exhibits significantly hindered electron transfer activity toward both electrodes and soluble acceptors (11, 14). As expected, this strain showed significantly attenuated electron transfer to PAAPEO-ITO (Figure 3c). Additionally, there was an order-of-magnitude decrease in the electron transfer rate constant compared to the wild-type strain (Table 1). It should be noted that statistical error associated with accumulated electron transfer is dominated by variation in *S. oneidensis* EET rates among biological replicates, not from error associated with spectra collection or fitting (Figure S10). The collected spectra are highly resolved, meaning the signal is stable over the collection time period of ∼1 minute and the fitting procedure consistently matches experimental data and converges to the same solution even for a range of randomly generated initial guesses (Table S6).

**Table 1.** Metrics obtained from HEDA fitting during EET from *S. oneidensis*. Data shown are results of the steady-state regime, between 1 and 8 h, and are the mean ± SD of *n* = 3 HEDA model fits of biological replicates.

| Sample | $e^-$ Transferred $\cdot \text{Cell}^{-1}$ | $k_{EET}$ ($e^- \cdot \text{cell}^{-1} \cdot \text{h}^{-1}$) | Current $\cdot \text{Cell}^{-1}$ (fA) |
| --- | --- | --- | --- |
| *S. oneidensis* MR-1<br>(+ 20 mM lactate) | $(1.3 \pm 0.2) \times 10^8$ | $(2.0 \pm 0.2) \times 10^7$ | $0.89 \pm 0.07$ |
| *S. oneidensis* MR-1<br>(– 20 mM lactate) | $(2.5 \pm 3.1) \times 10^7$ | $(4.8 \pm 2.4) \times 10^6$ | $0.21 \pm 0.11$ |
| *S. oneidensis*<br>$\Delta mtrC\Delta omcA\Delta mtrF$<br>(+ 20 mM lactate) | $(0.9 \pm 1.4) \times 10^7$ | $(1.2 \pm 1.2) \times 10^6$ | $0.05 \pm 0.05$ |

Although the physical dimensions of the nanocrystals (5.78 ± 0.64 nm radius, compared to cell length on the ∼1 µm scale) should prevent biofilm formation, we sought to ensure that EET was occurring from planktonic cultures. Therefore, we tested a biofilm-deficient knockout (*S. oneidensis* S2933, termed Δlysis-operon) (48). This strain did not exhibit significantly different current generation than wild-type *S. oneidensis*, indicating that electron transfer is occurring from planktonic cells (Figure 4). We repeated the wild-type sample as this experiment was performed with a newly synthesized batch of nanocrystals compared to Figure 3 (Note S3).

**Figure 4.**
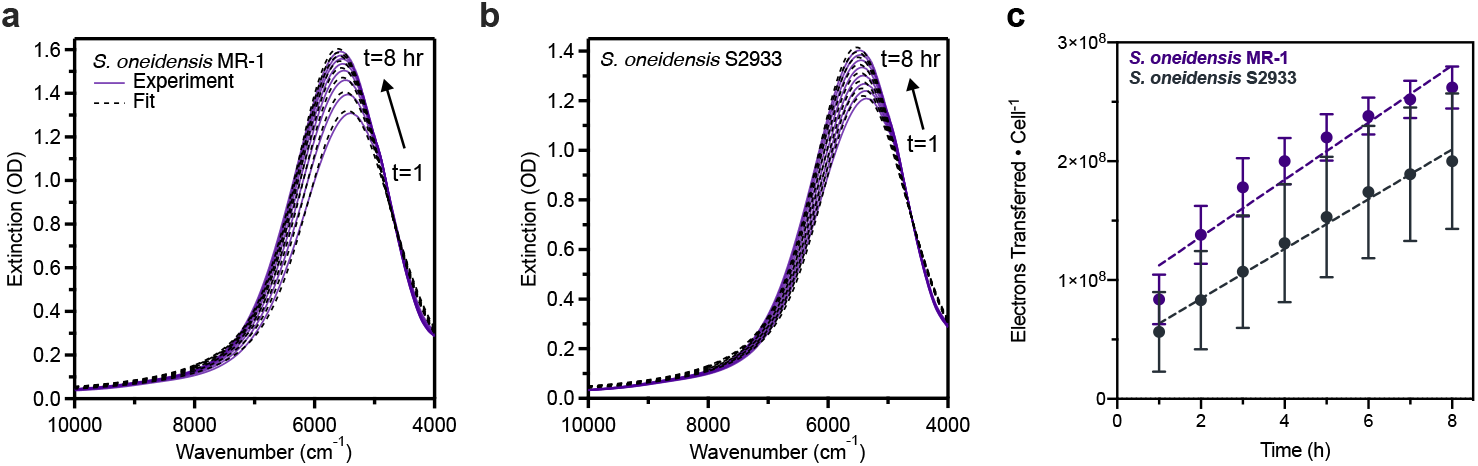
Planktonic cells are responsible for EET to PAAPEO-ITO. Extinction spectra for PAAPEO-ITO dispersions incubated with (a) *S. oneidensis* MR-1 and (b) and *S. oneidensis* S2933 (Δlysis-operon, a biofilm-deficient knockout) (48). (c) Steady-state EET (1–8 h) where dashed lines represent linear regression of the electrons transferred over time and *k*_*EET*_ is the slope of the regression (Table 1). For MR-1, *k*_*EET*_ = (2.4 ± 0.2) × 10^7^ e^−^ • cell^−1^ • h^−1^; for S2933, *k*_*EET*_ = (2.1 ± 0.4) × 10^7^ e^−^ • cell^−1^ • h^−1^. Data points shown are the mean ± SD of *n* = 3 HEDA model fits of biological replicates.

All fit results are reported in Tables S4-7. Overall, current quantified from the wild-type strain with 20 mM lactate is comparable to, but slightly lower than, results from other methods that have measured or extrapolated single cell electron flux from *S. oneidensis* to heterogeneous acceptors.

Previous studies found current generation from *S. oneidensis* can vary, from ∼2 fA • cell^−1^ in cell suspensions to soluble riboflavin, to ∼100-200 fA • cell^−1^ in single cells to electrodes (18, 23, 49). Methods measuring larger reduction rates have varied in experimental conditions, including extrapolating single cell EET from biofilms on electrodes or hematite nanoparticles (14, 18–19, 49–50). One primary difference in our study compared to these is the lack of cell immobilization on a surface. Cell motility may inhibit certain EET mechanisms such as flavin shuttling, which occurs primarily in attached cytochromes (19, 51). Therefore, it is possible that both physiological and experimental differences account for increased EET measurement in these platforms compared to ours. In addition, the necessity of deuterium in our buffer would lead to kinetic isotope effects on proton pumping and therefore electron transfer kinetics. Previous studies have determined that *S. oneidensis* can recover from this and reach deuterium efflux equilibrium (52); however, this may contribute to lower overall current generation measured in our system compared to others (Note S1). Finally, the polymer coating around the nanocrystals could increase the required distance for direct electron transfer, which would also reduce EET rate. Despite these challenges, our results indicate that the decreased rate does not inhibit us from differentiating between the EET rates for strains of varying metabolic activity and genotype. Overall, these results contribute to a quantitative understanding of EET from planktonic cell populations and suggest that simple spectroscopic techniques can be used as a screening tool to elucidate the electroactive capabilities of various species or strains.

### EET output from engineered *S. oneidensis* with variable *mtrC* expression can be quantitatively differentiated using PAAPEO-ITO

Synthetic biologists commonly use genetic circuits to control protein output, and emergent applications of EET will likely employ similar strategies. However, unlike fluorescence or secreted protein, it is difficult to directly quantify the relationship between EET gene expression and resultant electron flux, especially in planktonic cultures. Toward this goal, we used an *S. oneidensis* strain that tailors expression of a critical EET gene, *mtrC*, in response to inducing molecule concentration. Genetically engineered *S. oneidensis* Δ*mtrC*Δ*omcA*Δ*mtrF* + *mtrC* proportionally expresses *mtrC* in the presence of isopropyl ß-D-1-thiogalactopyranoside (IPTG) as an inducer (12, 46). Cultures of *S. oneidensis* Δ*mtrC*Δ*omcA*Δ*mtrF* + *mtrC* were pregrown anaerobically in varying concentrations of IPTG (ranging from 0 to 1000 µM), leading to differing expression levels of *mtrC* and resultant EET flux. Cultures were then washed and normalized to OD_600_ = 0.2 in a reaction mixture containing PAAPEO-ITO, 20 mM lactate, 25 µg/mL kanamycin for plasmid maintenance, and the corresponding concentration of inducing molecule. As outlined above, spectra were collected through 8 hours and fit with the HEDA model (Figure 5a–d). Fit results are reported in Table S8. As expected, the resultant electron transfer rate during steady-state was a function of IPTG concentration (Figure 5e), increasing about 5.5-fold from no induction/leaky expression (0 µM IPTG) to full induction (1000 µM IPTG). The consistent linear electron transfer rate across all engineered strains after 1 h again supports steady-state electron transfer. A wild-type (MR-1), empty vector control grown in 1000 µM IPTG also showed a high electron transfer rate, even higher than the wild-type without a plasmid, which could be attributed to the presence of antibiotic, physiological changes from plasmid maintenance, and/or an anaerobic pregrowth. In this sample, saturation effects were observed at 4 hours, so the rate constant was derived from only the preceding timepoints during steady-state electron transfer (Figures S11–12). It is notable that even the fully induced *mtrC* complement did not completely recapture the wild-type EET phenotype, exhibiting ∼4.5-fold less activity compared to the empty vector control. This may highlight the importance of other missing EET components in this strain, specifically *omcA* and *mtrF*, in binding and electron transfer to metal oxide substrates (53, 54). Tiered electron flux based on varying gene expression was also quantified as current output, ranging from 0.12 ± 0.04 (uninduced) to 0.68 ± 0.08 (fully induced) fA • cell^−1^ at steady-state (Table 2). The rate constants were fit with an activating Hill Function gene expression model as a guide to the eye; as expected, increasing concentrations of IPTG correspond to increasing values of *k*_*EET*_ (Figure 5f, Note S4). Together, these results exemplify the quantitative capabilities of our method for measuring electron flux after varying expression levels of a single EET gene. This deepens our understanding of the relationship between *mtrC* expression and EET output to heterogenous acceptors in planktonic cultures, and may be used to predict electron flux based on *mtrC* expression for a desired application.

**Table 2.** Metrics obtained from HEDA fitting during EET from engineered *S. oneidensis* MR-1 + *empty* and Δ*mtrC*Δ*omcA*Δ*mtrF* + *mtrC*. Data shown are results of the steady-state regime, between 1 and 8 h for all samples except for MR-1 + *empty*, which was between 1 and 3 h. Data shown are the mean ± SD of *n* = 3 HEDA model fits of biological replicates.

| Sample | e<sup>-</sup> Transferred • Cell<sup>-1</sup> | $k_{EET}$ (e<sup>-</sup> • cell<sup>-1</sup> • h<sup>-1</sup>) | Current • Cell<sup>-1</sup> (fA) |
| --- | --- | --- | --- |
| *S. oneidensis* MR-1<br>+ empty (1000 μM IPTG) | $(3.2 \pm 1.0) \times 10^8$ | $(6.2 \pm 1.3) \times 10^7$ | $2.76 \pm 0.60$ |
| *ΔmtrCΔomcAΔmtrF*<br>+ *mtrC* (1000 μM IPTG) | $(1.0 \pm 0.3) \times 10^8$ | $(1.5 \pm 0.2) \times 10^7$ | $0.68 \pm 0.08$ |
| *ΔmtrCΔomcAΔmtrF*<br>+ *mtrC* (100 μM IPTG) | $(8.3 \pm 1.9) \times 10^7$ | $(1.2 \pm 0.1) \times 10^7$ | $0.53 \pm 0.05$ |
| *ΔmtrCΔomcAΔmtrF*<br>+ *mtrC* (10 μM IPTG) | $(4.1 \pm 1.3) \times 10^7$ | $(5.6 \pm 1.1) \times 10^6$ | $0.25 \pm 0.05$ |
| *ΔmtrCΔomcAΔmtrF*<br>+ *mtrC* (0 μM IPTG) | $(1.9 \pm 1.3) \times 10^7$ | $(2.6 \pm 1.0) \times 10^6$ | $0.12 \pm 0.04$ |

**Figure 5.**
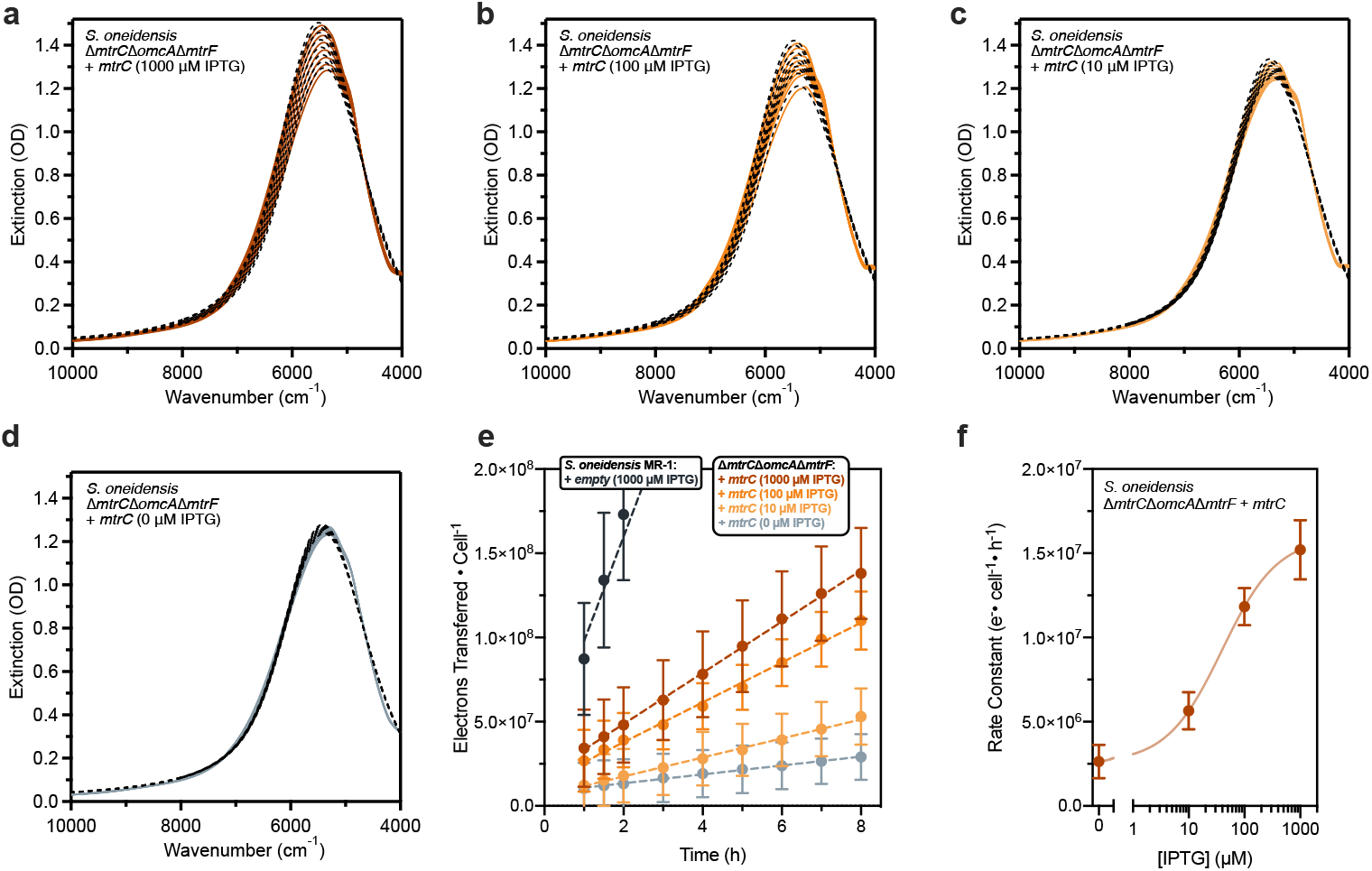
HEDA fitting of PAAPEO-ITO LSPR response during EET from engineered *S. oneidensis* reveals gradients in population current output as a result of differential gene expression. *In situ* extinction spectra and HEDA fits for PAAPEO-ITO during EET from (a–d) *S. oneidensis* Δ*mtrC*Δ*omcA*Δ*mtrF* + *mtrC* with (a) 1000, (b) 100, (c) 10, and (d) 0 µM inducing molecule (IPTG). (e) Steady-state EET (1–8 h) where dashed lines represent linear regression of the electrons transferred over time and *k*_*EET*_ is the slope of the regression (Table 2). Data points shown are the mean ± SD of *n* = 3 HEDA model fits of biological replicates. (f) Rate constants exhibit an activating Hill Function-type response to increasing IPTG concentration. Data points shown are the mean ± SD of *n* = 3 independent linear regressions of HEDA model fits of biological replicates.

## Conclusions

Together, our results demonstrate that the sensing capabilities of plasmonic ITO nanocrystals in microbial suspensions can expand the microbiologist toolkit for quantifying biological electron transfer. Using our platform, we successfully measured current generation on insoluble substrates from planktonic cultures, enabling differentiation of *S. oneidensis* cells of varying metabolic activity and genotype. In addition, we quantified changes in current generation by cells engineered to express varying levels of a critical EET gene, *mtrC*. This will enable fundamental studies of biological electron transfer metrics and resolution of EET gene expression dynamics. For example, PAAPEO-ITO biosensors can explore, in real time, how cells change electron flux in response to their environment, including the presence of a new electron acceptor or an inducing molecule. This non-destructive, *in situ* spectroscopic measurement could also be integrated with high-throughput technologies such as infrared microplate readers to screen genetic mutants or new bacterial strains based on electron transfer rate. In addition, infrared detection of EET enables simultaneous collection of visible wavelength metrics like growth rate, metabolic dyes, and fluorescent protein expression. We also envision PAAPEO-ITO could be used to monitor EET in visibly opaque environments, such as soil, but this will require more experiments to validate.

Our recent progress in both surface functionalization and spectroelectrochemical modeling have enabled this demonstration of PAAPEO-ITO as an accurate EET sensing platform; however, further advances in each of these areas will enable even more applications. For example, continuing to develop novel nanocrystal surface chemistry, such as functionalizing with bioactive polymers, could expand our capability to target specific cellular machinery, including surface proteins such as MtrC. In addition, standardizing the optical signatures of ITO nanocrystals with known surface potentials could allow both electron tracking and mapping of potential energy landscapes across the cell. Overall, our design highlights concurrent progress in nanocrystal biosensors and synthetic biology for EET, enabling more effective evaluation of critical biological processes.

## Materials and Methods

### Chemicals and reagents

Deuterium oxide (D_2_O, Sigma-Aldrich, 99.9 atom % D), sodium DL-lactate (NaC_3_H_5_O_3_, TCI, 60% in water), sodium fumarate (Na_2_C_4_H_2_O_4_, VWR, 98%), HEPES buffer solution (C_8_H_18_N_2_O_4_S, VWR, 1 M in water, pH = 7.3), potassium phosphate dibasic (K_2_HPO_4_, Sigma-Aldrich), potassium phosphate monobasic (KH_2_PO_4_, Sigma-Aldrich), sodium chloride (NaCl, VWR), ammonium sulfate ((NH_4_)_2_SO_4_, Fisher Scientific), magnesium (II) sulfate heptahydrate (MgSO_4_•7H_2_O, VWR), trace mineral supplement (ATCC), casamino acids (VWR), silicone oil (Alfa Aesar), N,N-dimethylformamide (DMF, Sigma-Aldrich, ≥99.8%), nitrosonium tetrafluoroborate (NOBF_4_, Sigma-Aldrich, 95%), ammonium cerium (IV) nitrate (Sigma-Aldrich, ≥98.5%), toluene (Fisher Scientific, ≥99.5%), tin (IV) acetate (Sigma Aldrich), indium (III) acetate (STREM 99.99%), oleic acid (Sigma Aldrich, technical grade, 90%), oleyl alcohol Sigma Aldrich, technical grade, 85%), hexane (Fisher Scientific, various methylpentanes 4.2%, ≥98.5%) isopropyl ß-D-1-thiogalactopyranoside (IPTG, Teknova), kanamycin sulfate (C_18_H_38_N_4_O_15_S, Growcells), nail polish (Electron Microscopy Sciences), BacLight Live/Dead Stain (Invitrogen) were all used as purchased unless otherwise noted. All media components were autoclaved or sterilized using 0.2 μm PES filters.

### Nanocrystal synthesis

Tin-doped indium oxide nanocrystals were synthesized using a slow-injection approach pioneered by the Hutchison group (39, 55). Indium acetate and tin acetate in a 9:1 molar ratio were dissolved in oleic acid at an overall concentration of 0.5 mmol metal per mL oleic acid. The solution was heated to 150°C and kept at that temperature for at least two hours to convert the metal-acetate complexes into the metal-oleate precursor. Using a syringe pump, the precursor was injected into a hot bath (290 °C) of oleyl alcohol at a rate of 0.3 mL/min. For this work, 4.5 mL of precursor was injected. The nanocrystals were collected from the reaction mixture and purified by flocculating with ethanol, centrifuging at 9000 rpm, and dispersing in hexane three consecutive times.

### Nanocrystal ligand stripping

NOBF_4_ was used to chemically remove the native organic ligands bound to the nanocrystal surface following a previously established method^31^. Briefly, 60 mg of NOBF_4_ were added to a two-phase mixture of DMF (2 mL) and ITO nanocrystal dispersion in hexane (∼50 mg/mL, 2 mL). The mixture was sonicated for 30 min to promote ligand removal and therefore the phase transfer of nanocrystals from hexane to DMF. After discarding the hexane layer, charge-stabilized ITO nanocrystals dispersed in DMF were purified by performing seven cycles of flocculation with toluene, centrifugation at 7500 rpm for 5 min, and redispersion in DMF.

### Nanocrystal polymer wrapping

PAA-mPEO_4_ random copolymer was synthesized (40, 41) and ITO nanocrystals were polymer-wrapped with PAA-mPEO_4_ following previously established methods (42). Briefly, a PAA-mPEO_4_ solution was prepared by dissolving 10 mg of polymer in 700 µL of DMF. Then, 300 µL of ITO nanocrystal dispersed in DMF (∼90 mg/mL) were added dropwise to the PAA-mPEO_4_ solution under gentle stirring. This nanocrystal-polymer mixture was stirred overnight and subsequently added dropwise into 19 mL of Milli-Q water while stirring. The aqueous mixture was stirred at 600 rpm for 48 hours. Polymer-wrapped ITO nanocrystals were recovered and purified by performing three cycles of spin dialysis (centrifugation at 4000 rpm for 10 min per cycle) using 50 kDa Millipore Amicon Ultra centrifugal tubes. The final dispersion in Milli-Q water was filtered through a 0.45 µm PVDF syringe filter (Acrodisc, Pall).

### Oxidation of polymer-wrapped nanocrystals

Polymer-wrapped ITO nanocrystals dispersed in Milli-Q water were chemically oxidized following a previously established method (36). Briefly, 120 µL of polymer-wrapped ITO (∼7 mg/mL) were mixed with 25 µL of an aqueous solution of ammonium cerium nitrate (25 mM). Instead of scaling up the volume of the reaction, the reagents were combined in multiple separate vials to achieve the desired final mass of oxidized and polymer-wrapped nanocrystals. The mixture was left to stand without stirring for 15 min and then purified by performing three cycles of spin dialysis (centrifugation at 4000 rpm for 10 min per cycle) using 50 kDa Millipore Amicon Ultra centrifugal tubes. The resulting oxidized and polymer-wrapped ITO nanocrystals were then dispersed in deuterated SBM + CAAs (Table S1) by performing a solvent exchange through three additional spin dialysis cycles with excess deuterated buffer instead of Milli-Q water (final concentration: 1.5 mg/mL).

### Nanocrystal Characterization

**a. Electron microscopy** ITO nanocrystals were imaged in bright-field scanning transmission electron microscopy mode at a 30 kV accelerating voltage on a Hitachi S5500 SEM/STEM instrument. The sample was drop-cast on Type-A ultrathin carbon copper TEM grids (Ted Pella, 01822, 400 mesh) from dilute ITO dispersions in hexane. **b. Small-angle X-ray scattering (SAXS)**. The size of as-synthesized ITO nanocrystals was determined by SAXS. A dilute dispersion (∼ 1 mg/mL), enclosed in a sealed glass capillary (Charles-Supper Company, Boron Rich, 1.5 mm diameter, 0.01 mm wall thickness), was measured in transmission configuration at a sample-detector distance of 1 m on a SAXSLAB Ganesha instrument using Cu K**α** radiation. Details of SAXS data analysis are included in the Supplementary Information. **c. Dynamic light scattering**. The hydrodynamic diameters of ligand-stripped, polymer-wrapped, and oxidized ITO nanocrystals were measured on a Malvern Zetasizer Nano ZS. Dilute ITO nanocrystals dispersions (∼1 mg/mL) were enclosed in disposable plastic micro cuvettes (ZEN0040, Malvern). **d. Thermogravimetric analysis**. PAA-mPEO_4_ adsorption on the ITO nanocrystal surface was quantified using a Mettler Toledo TGA 2. In a typical experiment, 100 µL of polymer-wrapped ITO nanocrystals dispersed in Milli-Q water (∼10 mg/mL) was added to a disposable aluminum crucible and dried under vacuum at room temperature for 24 hours. The sample was measured in dry synthetic air from 30 °C to 600 °C at a 5 °C/min ramp rate. **e. Fourier transform infrared spectroscopy**. Ligand-capped, ligand-stripped, and polymer-wrapped ITO nanocrystals were measured in transmission geometry with a Bruker Vertex 70 spectrometer. Samples were drop-cast from dilute nanocrystal dispersion (∼1 mg/mL) on undoped, double side polished silicon substrates (Virginia Semiconductor Inc.) **f. UV/Vis/NIR Spectroscopy**. The cuvettes were placed in an Agilent Cary series ultraviolet-visible-near infrared spectrophotometer in transmission mode for spectra collection. Spectra were collected from 3032-10000 cm^−1^ for a collection time of ∼1 min per spectrum. **g. Inductively coupled plasma-optical emission spectroscopy (ICP-OES)**. To quantify the volume fraction of nanocrystals in dispersion, as well as the doping percentage of tin atoms in the indium oxide lattice, we took a known volume of the dispersed nanocrystals, dried them into a pellet and dissolved them in aqua regia for two days. We then diluted the dissolved ions to 2% v/v acid and loaded the dilute acid into the Varian 720-ES ICP-OES to quantify the tin and indium concentrations. We calibrated the instrument for tin and indium using standards purchased from Sigma Aldrich (TraceCERT® 1000mg In/L nitric acid and 1000mg Sn/L nitric acid).

### Bacterial strains and culture

Bacterial strains and plasmids are listed in Table S3. Cultures were prepared as follows: bacterial stocks stored in 22.5% glycerol at −80 °C were streaked onto LB agar plates (for wild-type and knockout strains) or LB agar with 25 µg/mL kanamycin (for plasmid-harboring strains) and grown overnight at 30 °C for *Shewanella*, 37 °C for *E. coli*. Single colonies were isolated and inoculated into *Shewanella* Basal Medium (SBM) supplemented with 100 mM HEPES, 0.05% trace mineral supplement (Table S2), 0.05% casamino acids, and 20 mM sodium lactate (2.85 µL of 60% w/w sodium lactate per 1 mL culture) as the electron donor. Aerobic cultures were pregrown in 15 mL culture tubes at 30 °C and 250 rpm shaking. Anaerobic, plasmid-harboring cultures were pregrown using the same procedure outlined above, but in degassed growth medium in a humidified anaerobic chamber, supplemented with 40 mM sodium fumarate (40 µL/mL of a 1 M stock) as the electron acceptor, 25 µg/mL kanamycin, and varying concentrations of IPTG. Descriptions of the *mtrC* circuit and pCD24r1 plasmid, as well as the empty vector circuit and the pCD8 plasmid, have been published previously (12). Cultures were washed 3x after pregrowth using SBM supplemented with 0.5% casamino acids (degassed for anaerobic cultures). Before the final centrifugation, OD_600_ was measured using a NanoDrop 2000C spectrophotometer and the culture volume normalized such that the final OD_600_ in the reaction with PAAPEO-ITO would be 0.2.

### Viability assessments

All microscopy was performed using a Nikon Ti2 Eclipse inverted epifluorescence microscope. Cells assessed for viability by microscopy were incubated with PAAPEO-ITO for 2 h at 30 °C. The cells were centrifuged, and the nanocrystals were washed away using 0.85% NaCl solution. The cell suspensions were then incubated in the dark in the BacLight Live/Dead stain mix (1.5 µL/mL Syto9, 2.5 µL/mL propidium iodide in 0.85% NaCl solution) for 20 minutes. Stained cells were then washed 3x in 0.85% NaCl solution to remove unbound dye. Cell suspensions were loaded onto glass microscope slides and a no. 1 coverslip placed on top. Nail polish was used to seal the coverslip to the glass slide to prevent evaporative losses. Fluorescence for each stain (green for Syto9, red for propidium iodide) was measured using GFP and Texas Red excitation/emission filter cubes on a Nikon Ti2 Eclipse, as outlined previously (7). Live and dead populations were quantified using thresholding and masking in Fiji 1.0. For colony counting, cell samples before and after incubation with PAAPEO-ITO were diluted by factors of 10^5^–10^7^ in LB medium, and 100 µL of these dilutions was plated onto LB-agar. The viable population (CFU/mL) for each sample was quantified by averaging 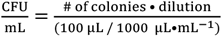 for each replicate.

### Electron transfer experiments and spectroscopy measurements

Bacteria and PAAPEO-ITO nanocrystals were mixed in a disposable centrifuge tube and immediately transferred to a Spectrocell near-IR quartz cuvette and the cuvette was sealed for the duration of the experiment. The cuvette was loaded into an Agilent Cary UV/vis/NIR spectrophotometer for spectra collection. On average, the first spectrum was taken ∼3 min after mixing. Thereafter, a spectrum was collected every 10 minutes for the first hour and then every hour until 8 hours. Three cuvettes were used to carry out triplicate experiments simultaneously, two with pathlengths of 1 mm and the third with pathlength 0.5 mm. All spectra were backgrounded to a cuvette filled solely with deuterated SBM buffer, without PAAPEO-ITO or bacteria.

## Author Contributions

A.J.G., S.L.G., C.A.S.C., D.J.M., and B.K.K. conceived the project; A.J.G., S.L.G., C.A.S.C., Y.W., and A.M.G. performed experiments; S.L.G., C.A.S.C., and Y.W. performed nanocrystal synthesis and characterization; A.J.G. designed biological experiments and performed microbial culture; A.J.G., S.L.G., C.A.S.C., D.J.M., and B.K.K. analyzed the results; A.J.G., S.L.G., C.A.S.C., D.J.M., and B.K.K. wrote the manuscript with input from all authors.

## Acknowledgements

The authors declare that they have no competing interests. All data needed to evaluate the conclusions in the paper are present in the paper and/or the Supplementary Materials. *S. oneidensis* Δ*mtrC*Δ*omcA*Δ*mtrF* was a generous gift from Prof. Jeffrey Gralnick (U. Minnesota), and *E. coli* MG1655 and *S. oneidensis* S2933 were generous gifts from Prof. Lydia Contreras (UT Austin). Christopher Dundas is thanked for his contribution of the engineered *S. oneidensis* strain. A.J.G. and S.L.G. were supported through National Science Foundation Graduate Research Fellowships (Program Award No. DGE-1610403). Portions of this work, including polymer synthesis and characterization by Swagat Sahu and Brett A. Helms, were carried out as a user project at the Molecular Foundry, which is supported by the Office of Science, Office of Basic Energy Sciences of the U.S. Department of Energy under contract no. DE-AC02-05CH11231. This research was primarily supported by the National Science Foundation through the Center for Dynamics and Control of Materials: an NSF Materials Research Science and Engineering Center under DMR-1720595. Additional research support was provided by the Welch Foundation (Grants F-1929 and F-1848), the National Institute of General Medical Sciences of the National Institutes of Health under Award Number R35GM133640 (B.K.K.), and an NSF CAREER Award (1944334, B.K.K.). The authors acknowledge use of shared research facilities supported in part by the Texas Materials Institute including the SAXS instrument acquired under an NSF MRI grant (CBET-1624659), the Center for Dynamics and Control of Materials: an NSF MRSEC (DMR-1720595), and the NSF National Nanotechnology Coordinated Infrastructure (ECCS-1542159). We gratefully acknowledge the use of facilities within the core microscopy lab of the Institute for Cellular and Molecular Biology, University of Texas at Austin.

## Data Deposition

Experimental data supporting the findings in this study are publicly available through the Texas Data Repository (doi: https://doi.org/10.18738/T8/OU8RNO).

