## Supplementary Information for "*In Situ* Optical Quantification of Extracellular Electron Transfer using Plasmonic Metal Oxide Nanocrystals"

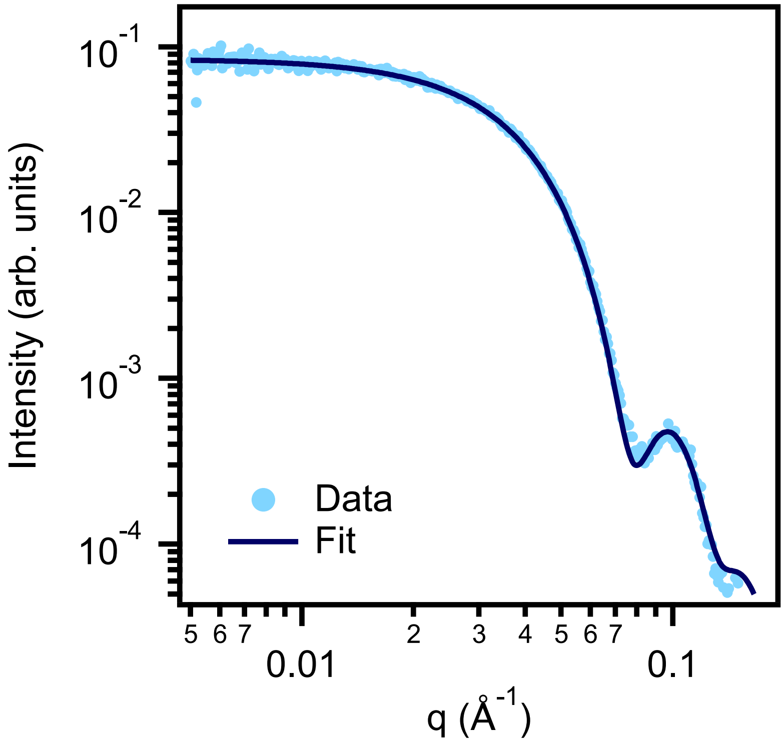


**Figure S1.** ITO nanocrystal sizing by small-angle X-ray scattering (SAXS). Experimental SAXS data of a dilute (~ 1 mg/mL) dispersion in hexane and spheroid model fit (solid line). Scattering data of a capillary containing neat hexane was collected for background subtraction. The scattering patterns were calibrated using a silver behenate standard[1] and were converted into 1D data by circular averaging using the Igor Pro-based Nika software for two-dimensional data reduction.[2] The Irena tool suite for modeling and analysis in Igor Pro was used for background subtraction and for fitting the nanocrystal form factor.[3]

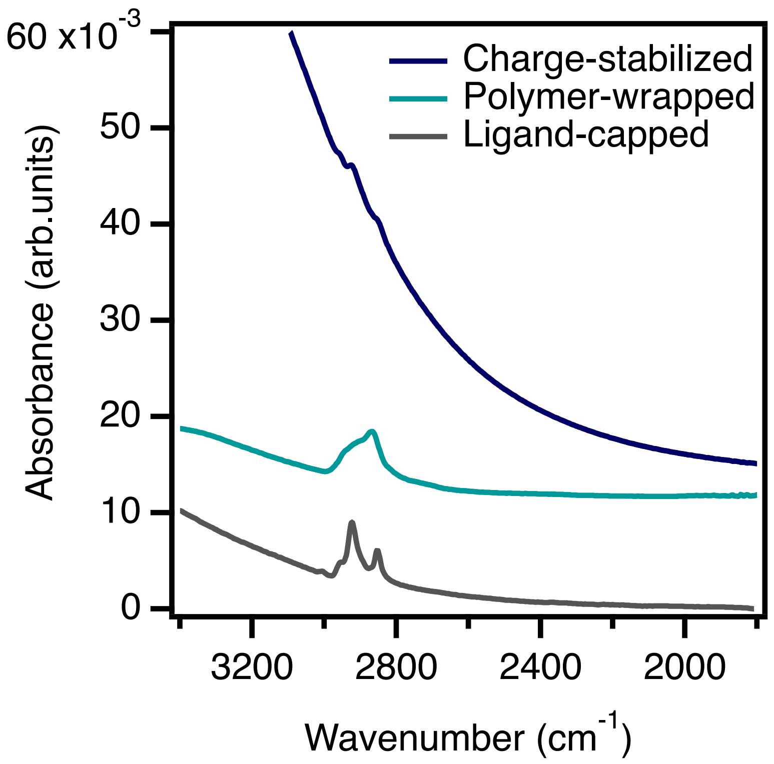


**Figure S2.** Fourier transform infrared spectroscopy spectra of as-synthesized ITO nanocrystals stabilized with organic ligands (ligand-capped, gray), ITO nanocrystals after ligand removal (charge-stabilized, blue), and ITO nanocrystals after functionalization with PAA-mPEO$_4$ (polymer-wrapped, teal). The spectra of ligand-capped and polymer-wrapped ITO nanocrystals exhibit peaks in the 2800–3000 cm$^{-1}$ region characteristic of C–H stretches from oleate ligands and the PAA-mPEO$_4$ backbone,[4–6] respectively. The absence of these peaks in the spectrum of charge-stabilized ITO nanocrystals indicates effective ligand removal.

**Table S1.** Ingredients in *Shewanella* Basal Medium (SBM). Growth media was supplemented with casamino acids and Wolfe's mineral solution, whereas PAAPEO-ITO reaction media was only supplemented with casamino acids, and ddH$_2$O was replaced with heavy water (D$_2$O).

| Ingredient | Quantity for 1L of 1x SBM |
|---|---|
| $K_2HPO_4$ | 225 mg |
| $KH_2PO_4$ | 225 mg |
| NaCl | 460 mg |
| $(NH_4)_2SO_4$ | 1.703 mL of 1 M stock |
| $MgSO_4 \cdot 7H_2O$ | 0.475 mL of 1 M stock |
| HEPES | 100 mL of 1 M stock |
| Casamino acids | 5 mL of 10% stock |
| Wolfe's mineral solution | 5 mL of 200x stock |
| ddH$_2$O | Up to 1 L, adjust to pH = 7.2 |

**Table S2**. Ingredients in Wolfe's mineral solution, adapted from ATCC recipe.

| Reagent | Quantity in 1L of 200X Stock |
|---|---|
| EDTA | 0.5 g (2.69 mL of 0.5 M stock) |
| $MgSO_4{\cdot}7H_2O$ | 3.0 g |
| $MnSO_4{\cdot}H_2O$ | 0.5 g |
| NaCl | 1.0 g |
| $FeSO_4{\cdot}7H_2O$ | 0.1 g |
| $Co(NO_3)_2{\cdot}6H_2O$ | 0.1 g |
| $CaCl_2$ | 0.9 mL from 1 M stock |
| $ZnSO_4{\cdot}7H_2O$ | 0.1 g |
| $CuSO_4{\cdot}5H_2O$ | 10 mg |
| $AlK(SO_4)_2$ | 10 mg |
| $H_3BO_3$ | 10 mg |
| $Na_2MoO4{\cdot}2H_2O$ | 10 mg |
| $Na_2SeO_3$ | 1 mg |
| $Na_2WO_4{\cdot}2H_2O$ | 10 mg |
| $NiCl_2{\cdot}6H_2O$ | 20 mg |
| $ddH_2O$ | 1 L |

**Note S1: Considerations for a deuterated buffer**

Because of the strong extinction of water near the LSPR of PAAPEO-ITO, optical quantification of EET necessitated the use of a deuterated buffer for *in situ* optical measurements, spectral fitting, and subsequent analysis of EET kinetics. The kinetic isotope effect (KIE) of deuterium in bacteria is a well-studied phenomenon that can strongly influence bacterial physiology and intracellular reaction rates. Okamoto and coworkers have demonstrated this phenomenon using *S. oneidensis* biofilms on an ITO electrode, and noted an initial decrease in electron transfer kinetics upon addition of $D_2O$.[7] In general, however, EET rate recovered on the order of minutes. While this effect may contribute to the initial variability in electron transfer rate in our system, the bacteria seemingly adjust and reach a deuterium efflux equilibrium, as supported by consistent steady-state electron transfer after 1 h and minimal effect on cell viability. How the KIE changes in planktonic vs. biofilm EET, as well as deuterium's effect on proton efflux mechanisms over longer periods in planktonic culture, require further investigation. Future work exploring alternative plasmonic metal oxide nanocrystals could circumvent the need for deuterated media. For example, rhenium oxide,[8] tungsten oxide,[9,10] and fluorine, tin co-doped indium oxide[11] all exhibit $\omega_{LSPR}$ at higher energy than ITO. With further dopant, size, and shape engineering, these materials hold promise for quantification of plasmonic responses within the water-transparent, near-IR region.

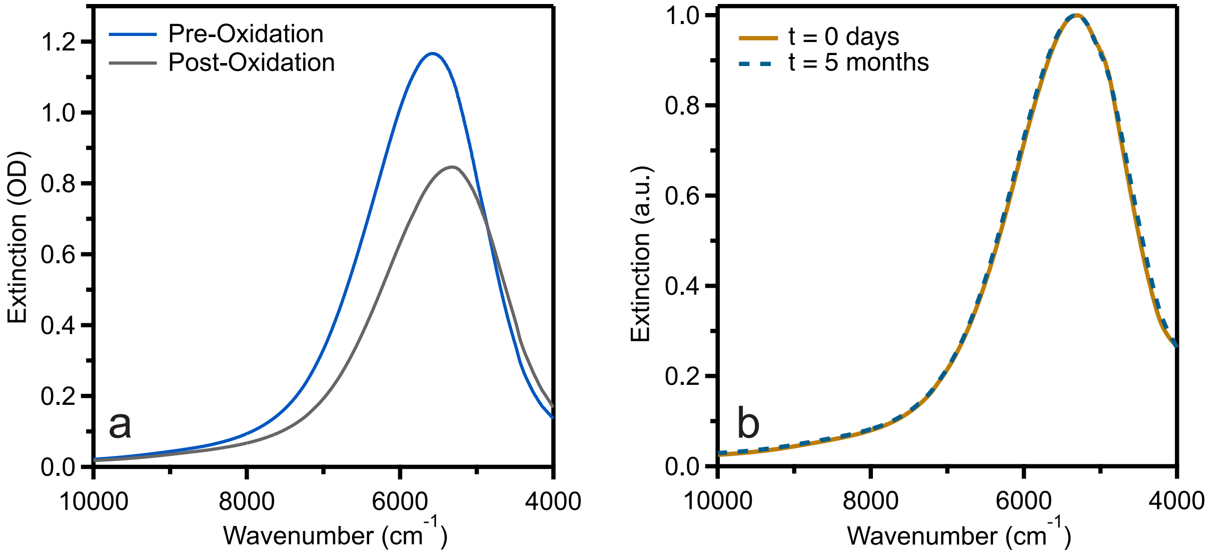


**Figure S3.** Oxidized nanocrystals remain stable in deuterated SBM. (a) Extinction spectra for PAAPEO-ITO dispersions in $D_2O$ before (pre-oxidation) and after (post-oxidation) oxidation with ammonium cerium nitrate. (b) Normalized extinction spectra immediately after oxidation and 5 months after oxidation indicate long-term colloidal stability.

**Table S3.** Bacterial strains and plasmids used in this study.

| Strain or plasmid | Description/Genotype | Reference or source |
|---|---|---|
| ***S. oneidensis* Strains** | | |
| MR-1 | MR-1 (ATCC700550), wild-type strain | American-Type Culture Collection |
| JG596 | Lacks out membrane cytochromes MtrC, OmcA, and MtrF; Δ*mtrC*Δ*omcA*Δ*mtrF* | [12] |
| MR-1 + pCD8 | Wild-type with an empty vector on the LacI repressed circuit | [13, 14] |
| JG596 + pCD24r1 | JG596 with a LacI repressed *mtrC* circuit | [13, 14] |
| S2933 | Biofilm-deficient Δlysis-operon strain | [15]; Lydia Contreras, U. of Texas at Austin |
| ***E. coli* Strains** | | |
| MG1655 | Wild-type strain | Lydia Contreras, U. of Texas at Austin |
| **Plasmids** | | |
| pCD8 | Empty vector antibiotic/inducer control | [13, 14] |
| pCD24r1 | *mtrC* expression vector, $P_{tac}$/*lacI* repressor cassette, $kan^R$ resistance (see ref) | [13, 14] |

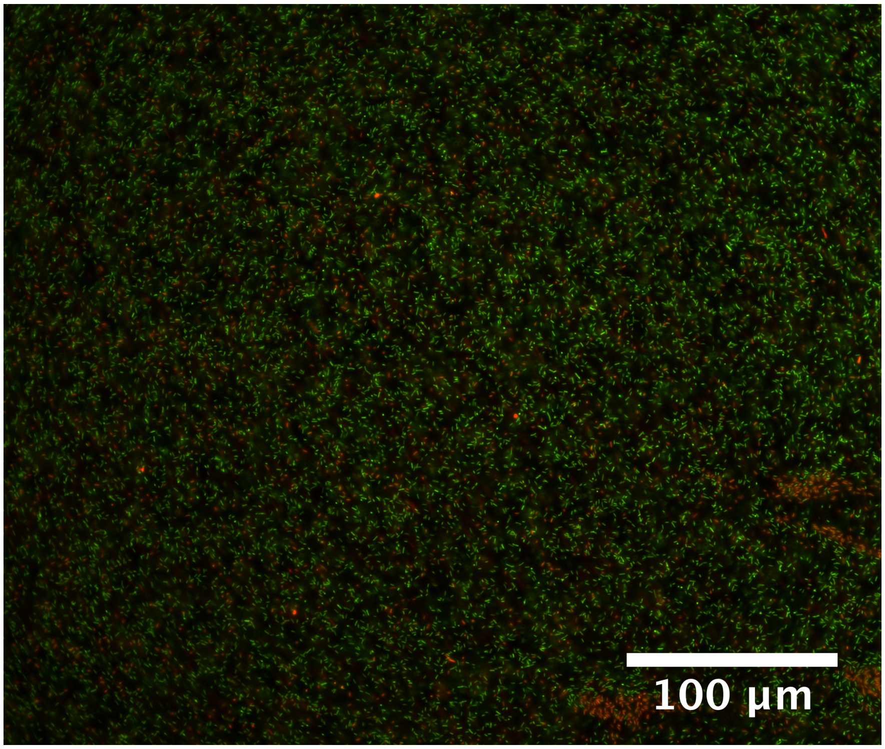


**Figure S4.** *S. oneidensis* remain viable in the presence of ~1 mg/mL PAAPEO-ITO. Cell viability (86.4 ± 2.24%) was quantified with the BacLight Live/Dead Stain (Invitrogen), using the GFP (live) and Texas Red (dead) fluorescence channels on a Nikon Ti2 Eclipse epifluorescence microscope.

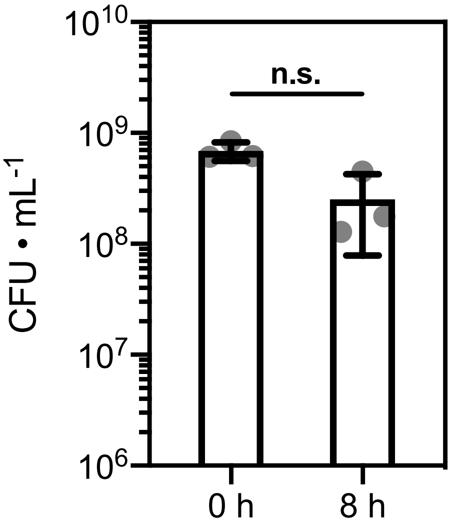


**Figure S5.** *S. oneidensis* viability is not substantially impacted during steady-state PAAPEO-ITO reduction. Colony counting of aerobically pregrown *S. oneidensis* MR-1 pre- and post-incubation with PAAPEO-ITO enabled population-average determination of current output from single cells. Data shown are mean ± S.D. of $n$ = 3 biological replicates.

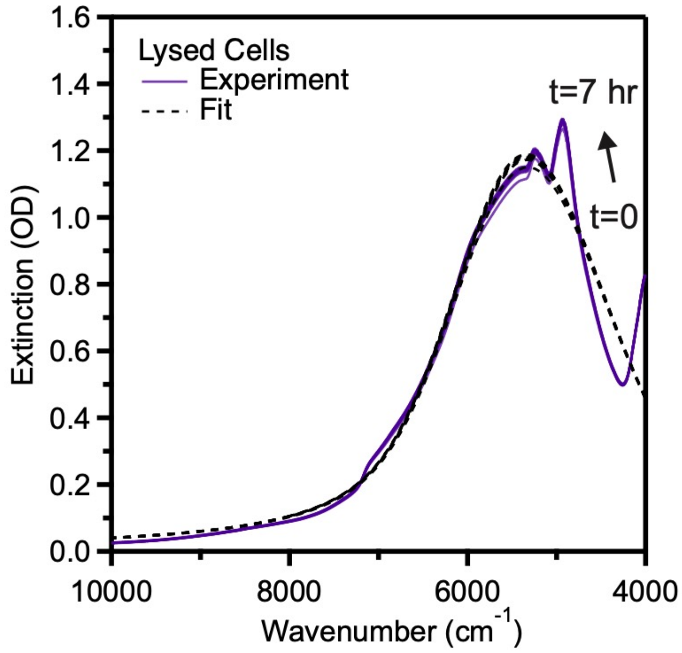


**Figure S6.** Cell lysate leads to minimal nonspecific reduction of PAAPEO-ITO. Extinction spectra for PAAPEO-ITO dispersions incubated with lysed *S. oneidensis* MR-1. Extraneous peaks at ~4800 $cm^{-1}$ are likely due to excess water required to prepare lysed cell samples.

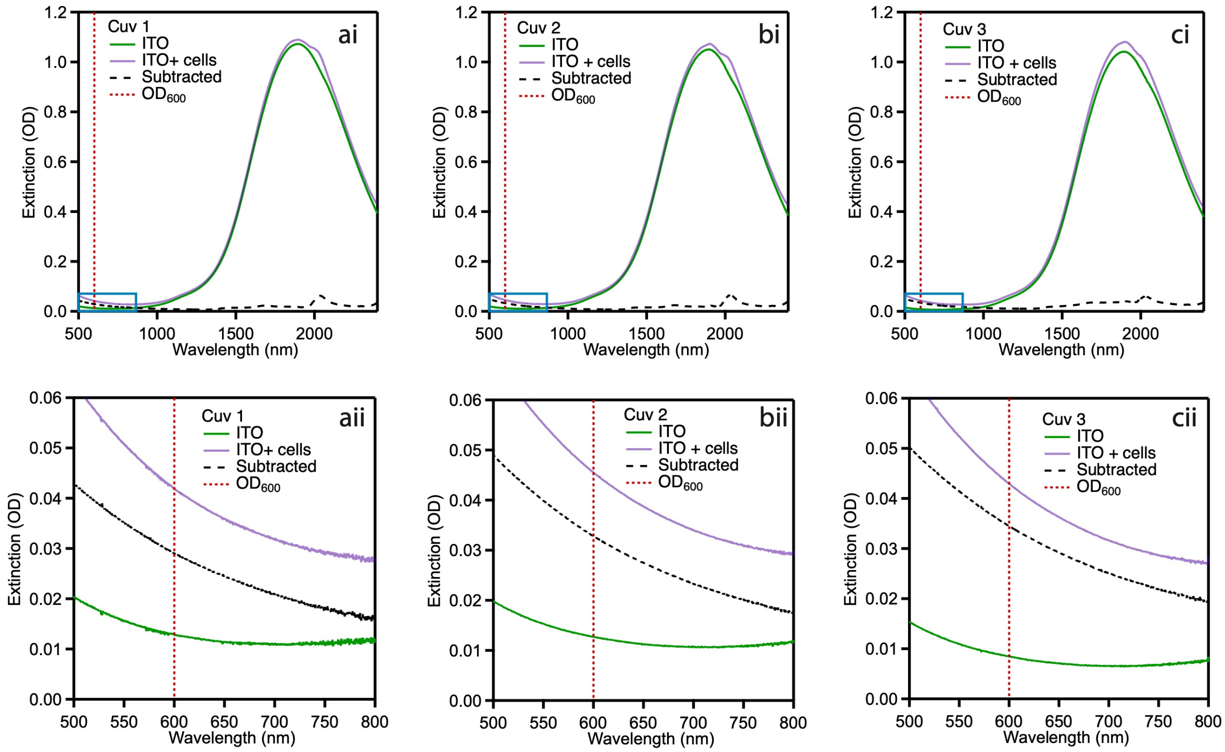


**Figure S7.** Simultaneous visible and infrared detection. *S. oneidensis* were diluted to $OD_{600}$ = 0.2 as measured by NanoDrop and then mixed with PAAPEO-ITO. Because the pathlength of the NanoDrop is 10 mm, we expect to measure $OD_{600}$ = 0.02 for 1 mm pathlength cuvettes. (ai, bi, ci) The full collected spectrum of the dispersed PAAPEO-ITO (green) is subtracted from the ITO + *S. oneidensis* mixture (purple) to result in the subtracted spectrum that is attributed to scattering from only the cells (black dashed line). The blue box denotes the boundary of the zoomed in graphs of each triplicate (aii, bii, cii). Our measurements were consistent, showing $OD_{600}$ = 0.030, 0.033, 0.034 for replicates 1, 2, and 3, respectively. This is ~1.5x the value as measured by NanoDrop, which could be due to pipetting error or cell growth during oxygen exposure. Data shown are individually collected spectra from $n$ = 3 biological replicates.

**Note S2: HEDA fitting.**

The HEDA model employed in this work was identical to that described in prior work[16]. In brief, HEDA uses a least-squares fit function in MATLAB to minimize the error between the experimental and fitted spectrum. Given the average and standard deviation of the particle diameters acquired from SAXS, as well as the concentration of nanocrystals in dispersion, $C_{NC}$, the HEDA model fits for the average free electron concentration, $\mu_{n_e}$, the standard deviation in free electron concentration as it varies amongst nanocrystals in the ensemble, $\sigma_{n_e}$, volume fraction of the plasmonic core $F_{v,core}$, and the mean free path of an electron. We do not expect the mean free path of an electron to change much during an electron transfer experiment; therefore, we decided to fix this value to reduce the number of overall fit parameters. To acquire the mean free path, we fit six test spectra (all from oxidized PAAPEO-ITO before mixing with bacteria) and found the average mean free path to be 14 nm.

To calculate the number of electrons transferred per cell we first calculated the number of electrons per PAAPEO-ITO, $N_{NC}$. Given the extracted fit parameters for average electron concentration, $\mu_{n_e}$, and plasmonic core volume fraction $F_{v,core}$, we used Equation S1 to calculate $N_{NC}$, where $V$ is the average NC volume.

$$N_{NC} = \mu_{n_e} * V * F_{v,core} \quad \text{Equation S1}$$

We calculated the change in electrons per NC by finding the difference between $N_{NC}$ at two time points $f$ and $i$.

$$\Delta N_{NC} = N_{NC,f} - N_{NC,i} \quad \text{Equation S2}$$

Then, knowing both the concentration of NCs $C_{NC}$ and cells $C_{cell}$ in solution, we converted the change in electrons per NC to electrons transferred per cell between time points $f$ and $i$.

$$\Delta N_{cell} = \Delta N_{NC} * C_{cell}/C_{NC} \quad \text{Equation S3}$$

**Quantification of nanocrystal concentration, $C_{NC}$:**
After quantifying ppm $In^{3+}$ and $Sn^{4+}$ through ICP-OES (See Materials and Methods), we used the molar mass of $Sn_xIn_{2-x}O_3$ as well as the bulk density of indium oxide (7.14 g/mL) to calculate the concentration of nanocrystals in the deuterated buffer.

**Quantification of cell concentration, $C_{cell}$:**
For cell concentration, the approximate CFU/mL at the inoculating $OD_{600}$ was used (~$10^8$ CFU/mL).

**Statistical analysis and propagation of error.**
We ran each experiment for *n* = 3 biological replicates and collected spectra at identical time points. We extracted the average value and standard deviation of the fit parameters among the three experiments at each time point. From those values, we calculated the average values for $N_{NC}$, $\Delta N_{NC}$, and $\Delta N_{cell}$ as described in Equations S1–S3. We calculated the standard deviations for those values with the following equations for propagation of error:

$$\frac{\sigma_{N_{NC}}}{N_{NC}} = \sqrt{\left(\frac{\sigma_{n_e}}{\mu_{n_e}}\right)^2 + \left(\frac{\sigma_{F_{v,core}}}{F_{v,core}}\right)^2} \quad \text{Equation S4}$$

$$\sigma_{\Delta N_{NC}} = \sqrt{\sigma_{\mathrm{NC,f}}{}^2 + \sigma_{\mathrm{NC,i}}{}^2} \quad \text{Equation S5}$$

$$\sigma_{\Delta N_{cell}} = \sigma_{\Delta N_{NC}} * C_{cell}/C_{NC} \quad \text{Equation S6}$$

Unless otherwise noted, data are reported as mean ± SD of *n* = 3 biological replicates.

**Table S4.** HEDA fit results and calculated values quantifying electron transfer extracted from spectra plotted in Figure S6. Variables are defined and the standard deviation in each parameter is calculated as described in Note S2.

**Strain:** Lysed *S. oneidensis* MR-1

| | Fit Results | | | Calculated Parameters | | |
|---|---|---|---|---|---|---|
| Time (min) | $\mu_{n_e}$ (x $10^{20}$ $cm^3$) | $\sigma_{n_e}$ (x $10^{20}$ $cm^3$) | $F_{v,core}$ | $N_{NC}$ | $\Delta N_{NC}$ | $\Delta N_{\boldsymbol{cell}}$ (x $10^7$) |
| 0 | 9.5 ± 0.04 | 2.8 ± 0.03 | 0.685 ± 0.010 | 526 ± 8 | 0 | 0 |
| 60 | 9.5 ± 0.04 | 2.8 ± 0.03 | 0.694 ± 0.010 | 534 ± 8 | 7 ± 11 | 1.3 ± 2.0 |
| 120 | 9.5 ± 0.04 | 2.8 ± 0.02 | 0.689 ± 0.008 | 532 ± 7 | 6 ± 11 | 1.0 ± 1.8 |
| 180 | 9.5 ± 0.05 | 2.8 ± 0.02 | 0.691 ± 0.010 | 533 ± 8 | 7 ± 12 | 1.2 ± 2.0 |
| 240 | 9.5 ± 0.05 | 2.8 ± 0.03 | 0.693 ± 0.009 | 535 ± 7 | 8 ± 11 | 1.4 ± 1.9 |
| 300 | 9.5 ± 0.05 | 2.8 ± 0.03 | 0.696 ± 0.010 | 536 ± 8 | 10 ± 12 | 1.7 ± 2.0 |
| 360 | 9.5 ± 0.04 | 2.8 ± 0.03 | 0.701 ± 0.009 | 539 ± 7 | 13 ± 11 | 2.2 ± 1.9 |
| 450 | 9.5 ± 0.05 | 2.8 ± 0.03 | 0.703 ± 0.012 | 541 ± 10 | 14 ± 13 | 2.5 ± 2.2 |

**Table S5.** HEDA fit results and calculated values quantifying electron transfer extracted from spectra plotted in main text Figure 2. Variables are defined and the standard deviation in each parameter is calculated as described in Note S2.

**Strain:** *S. oneidensis* MR-1 + lactate

| | Fit Results | | | Calculated Parameters | | |
|---|---|---|---|---|---|---|
| Time (min) | $\mu_{n_e}$ (x $10^{20}$ $cm^3$) | $\sigma_{n_e}$ (x $10^{20}$ $cm^3$) | $F_{v,core}$ | $N_{NC}$ | $\Delta N_{NC}$ | $\Delta N_{\boldsymbol{cell}}$ (x $10^7$) |
| 0 | 11.3 ± 0.05 | 2.2 ± 0.01 | 0.512 ± 0.013 | 470 ± 12 | 0 | 0 |
| 60 | 11.4 ± 0.04 | 2.2 ± 0.01 | 0.543 ± 0.002 | 500 ± 3 | 30 ± 12 | 4.1 ± 1.7 |
| 90 | 11.4 ± 0.07 | 2.2 ± 0.01 | 0.546 ± 0.006 | 504 ± 6 | 34 ± 14 | 4.7 ± 1.9 |
| 120 | 11.5 ± 0.09 | 2.1 ± 0.01 | 0.551 ± 0.004 | 510 ± 5 | 41 ± 13 | 5.6 ± 1.8 |
| 180 | 11.5 ± 0.11 | 2.1 ± 0.01 | 0.566 ± 0.004 | 528 ± 6 | 58 ± 14 | 8.0 ± 1.9 |
| 240 | 11.6 ± 0.14 | 2.1 ± 0.01 | 0.584 ± 0.010 | 547 ± 12 | 77 ± 17 | 10.6 ± 2.3 |
| 300 | 11.6 ± 0.17 | 2.1 ± 0.01 | 0.598 ± 0.011 | 562 ± 13 | 92 ± 18 | 12.8 ± 2.5 |
| 360 | 11.7 ± 0.17 | 2.1 ± 0.01 | 0.609 ± 0.008 | 574 ± 11 | 104 ± 16 | 14.4 ± 2.3 |
| 420 | 11.6 ± 0.12 | 2.1 ± 0.01 | 0.626 ± 0.018 | 589 ± 18 | 119 ± 21 | 16.5 ± 3.0 |
| 480 | 11.7 ± 0.16 | 2.1 ± 0.00 | 0.629 ± 0.015 | 593 ± 16 | 123 ± 20 | 17.1 ± 2.8 |

**Strain:** *S. oneidensis* MR-1 - lactate

| | Fit Results | | | Calculated Parameters | | |
|---|---|---|---|---|---|---|
| Time (min) | $\mu_{n_e}$ (x $10^{20}$ $cm^3$) | $\sigma_{n_e}$ (x $10^{20}$ $cm^3$) | $F_{v,core}$ | $N_{NC}$ | $\Delta N_{NC}$ | $\Delta N_{\mathbf{cell}}$ (x $10^7$) |
| 0 | 11.4 ± 0.10 | 2.2 ± 0.01 | 0.503 ± 0.016 | 463 ± 15 | 0 | 0 |
| 60 | 11.4 ± 0.10 | 2.2 ± 0.02 | 0.502 ± 0.016 | 463 ± 15 | 0 ± 21 | 0.0 ± 2.9 |
| 90 | 11.4 ± 0.11 | 2.2 ± 0.02 | 0.504 ± 0.016 | 465 ± 16 | 1 ± 22 | 0.2 ± 3.0 |
| 120 | 11.4 ± 0.10 | 2.2 ± 0.02 | 0.502 ± 0.015 | 464 ± 14 | 1 ± 21 | 0.0 ± 2.9 |
| 180 | 11.4 ± 0.11 | 2.2 ± 0.02 | 0.503 ± 0.016 | 464 ± 16 | 1 ± 22 | 0.1 ± 3.0 |
| 240 | 11.4 ± 0.10 | 2.2 ± 0.02 | 0.507 ± 0.021 | 468 ± 20 | 5 ± 25 | 0.7 ± 3.4 |
| 300 | 11.4 ± 0.10 | 2.2 ± 0.01 | 0.521 ± 0.026 | 479 ± 24 | 16 ± 29 | 2.2 ± 4.0 |
| 360 | 11.4 ± 0.09 | 2.2 ± 0.01 | 0.520 ± 0.023 | 480 ± 21 | 17 ± 26 | 2.4 ± 3.6 |
| 420 | 11.4 ± 0.13 | 2.2 ± 0.01 | 0.527 ± 0.028 | 486 ± 26 | 23 ± 30 | 3.1 ± 4.2 |
| 480 | 11.4 ± 0.06 | 2.2 ± 0.01 | 0.520 ± 0.017 | 482 ± 16 | 18 ± 22 | 2.6 ± 3.1 |

**Strain:** *E. coli* MG1655

| | Fit Results | | | Calculated Parameters | | |
|---|---|---|---|---|---|---|
| Time (min) | $\mu_{n_e}$ (x $10^{20}$ $cm^3$) | $\sigma_{n_e}$ (x $10^{20}$ $cm^3$) | $F_{v,core}$ | $N_{NC}$ | $\Delta N_{NC}$ | $\Delta N_{\mathbf{cell}}$ (x $10^7$) |
| 0 | 10.7 ± 0.09 | 2.3 ± 0.04 | 0.567 ± 0.006 | 491 ± 7 | 0 | 0 |
| 60 | 10.7 ± 0.09 | 2.3 ± 0.04 | 0.571 ± 0.006 | 494 ± 7 | 3 ± 9 | 0.45 ± 1.6 |
| 120 | 10.7 ± 0.09 | 2.3 ± 0.04 | 0.571 ± 0.007 | 494 ± 7 | 3 ± 10 | 0.54 ± 1.7 |
| 180 | 10.7 ± 0.09 | 2.3 ± 0.05 | 0.573 ± 0.006 | 496 ± 6 | 4 ± 9 | 0.73 ± 1.6 |
| 240 | 10.7 ± 0.08 | 2.3 ± 0.04 | 0.576 ± 0.005 | 498 ± 5 | 6 ± 9 | 1.1 ± 1.5 |
| 300 | 10.7 ± 0.09 | 2.3 ± 0.04 | 0.581 ± 0.006 | 501 ± 7 | 10 ± 10 | 1.6 ± 1.7 |
| 360 | 10.6 ± 0.09 | 2.4 ± 0.04 | 0.587 ± 0.007 | 505 ± 7 | 13 ± 10 | 2.3 ± 1.7 |
| 420 | 10.6 ± 0.08 | 2.4 ± 0.04 | 0.589 ± 0.006 | 506 ± 6 | 15 ± 9 | 2.6 ± 1.6 |

**Strain:** Δ*mtrC*Δ*omcA*Δ*mtrF* + lactate

| | Fit Results | | | Calculated Parameters | | |
|---|---|---|---|---|---|---|
| Time (min) | $\mu_{n_e}$ (x $10^{20}$ $cm^3$) | $\sigma_{n_e}$ (x $10^{20}$ $cm^3$) | $F_{v,core}$ | $N_{NC}$ | $\Delta N_{NC}$ | $\Delta N_{\boldsymbol{cell}}$ (x $10^7$) |
| 0 | 10.9 ± 0.10 | 2.3 ± 0.03 | 0.565 ± 0.012 | 500 ± 11 | 0 | 0 |
| 60 | 11.0 ± 0.06 | 2.2 ± 0.01 | 0.567 ± 0.007 | 503 ± 6 | 3 ± 13 | 0.3 ± 1.7 |
| 90 | 11.0 ± 0.07 | 2.2 ± 0.02 | 0.568 ± 0.007 | 504 ± 7 | 4 ± 13 | 0.4 ± 1.7 |
| 120 | 11.0 ± 0.08 | 2.2 ± 0.01 | 0.568 ± 0.010 | 504 ± 9 | 4 ± 15 | 0.5 ± 1.7 |
| 180 | 11.0 ± 0.08 | 2.2 ± 0.01 | 0.568 ± 0.009 | 504 ± 9 | 4 ± 14 | 0.5 ± 1.7 |
| 240 | 11.0 ± 0.10 | 2.2 ± 0.02 | 0.567 ± 0.011 | 505 ± 11 | 4 ± 16 | 0.5 ± 1.7 |
| 300 | 11.0 ± 0.10 | 2.2 ± 0.02 | 0.568 ± 0.011 | 506 ± 11 | 6 ± 16 | 0.7 ± 1.7 |
| 360 | 11.0 ± 0.07 | 2.2 ± 0.02 | 0.572 ± 0.009 | 508 ± 9 | 8 ± 14 | 1.0 ± 1.7 |
| 420 | 11.0 ± 0.07 | 2.2 ± 0.01 | 0.573 ± 0.010 | 509 ± 9 | 9 ± 15 | 1.1 ± 1.7 |
| 480 | 11.0 ± 0.07 | 2.2 ± 0.02 | 0.574 ± 0.011 | 511 ± 10 | 10 ± 15 | 1.2 ± 1.7 |

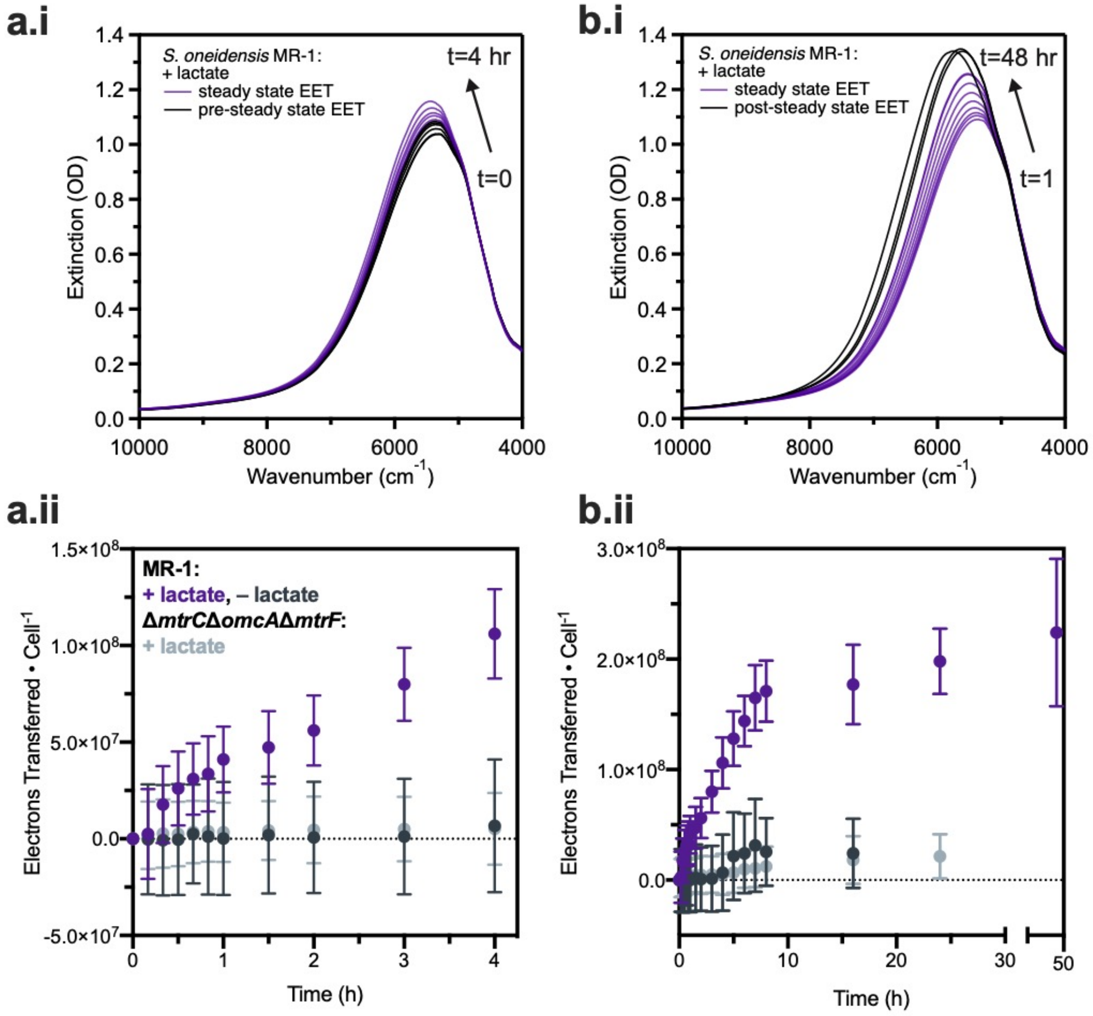


**Figure S8**. Time-resolved spectroscopic measurement of PAAPEO-ITO during EET from *S. oneidensis* MR-1 demonstrates steady-state behavior. (a) The early domain (≤1 h) shows variability due to aerobic respiration, introduction of a deuterated solvent, and cell stress. This regime includes a lag phase (0–20 min), followed by a sharp increase in electron transfer (20–60 min). This phenomenon is illustrated by (a.i) PAAPEO-ITO extinction spectra as well as (a.ii) cumulative electrons transferred per cell. (b) The early domain is followed by steady-state electron transfer (1–8 h). The later domain (≥8 h) shows effects of nanocrystal saturation and precipitation, as demonstrated by (b.i) PAAPEO-ITO extinction spectra as well as (b.ii) cumulative electrons transferred per cell. The domain between 1 and 8 h consistently showed steady-state behavior, and thus was used for fitting analysis. Data shown are mean ± S.D. of $n$ = 3 biological replicates.

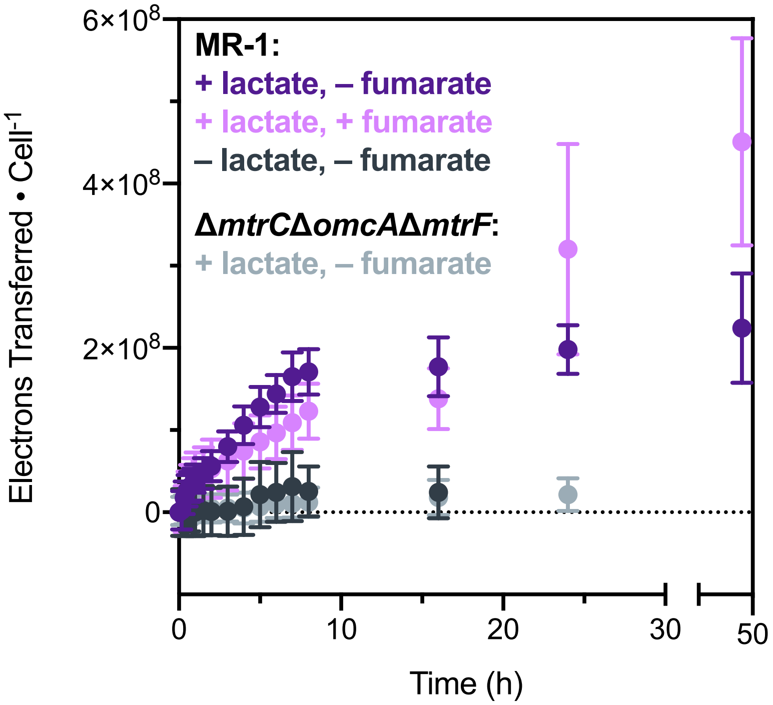


**Figure S9.** Additional electron acceptor, such as fumarate, enables sustained PAAPEO-ITO reduction for longer times. Due to potential effects of nanocrystal saturation and competition for electrons between PAAPEO-ITO and fumarate, fumarate was not included in future experiments. Data for all other samples are as shown in Figure S7. Data shown are mean ± S.D. of $n$ = 3 biological replicates.

**Note S3: Effect of nanocrystal concentration**

Both the concentration of nanocrystals in solution and the nanocrystal-to-cell ratio are important parameters that could be optimized for future sensing applications. We anecdotally observed slightly higher electron transfer rates with higher nanocrystal concentration. This result, combined with the relatively dilute nanocrystal concentration in these experiments, leads us to believe that electron transfer in our system is diffusion-limited. Hence, the rate of metabolic electron flux from *S. oneidensis* could reach higher values than those reported here. Still, the observed trends are expected to hold since each group of experiments conducted employed a single nanocrystal batch of constant concentration. A systematic study is required to investigate the effects of diffusion limitation and nanocrystal concentration on electron transfer rates.

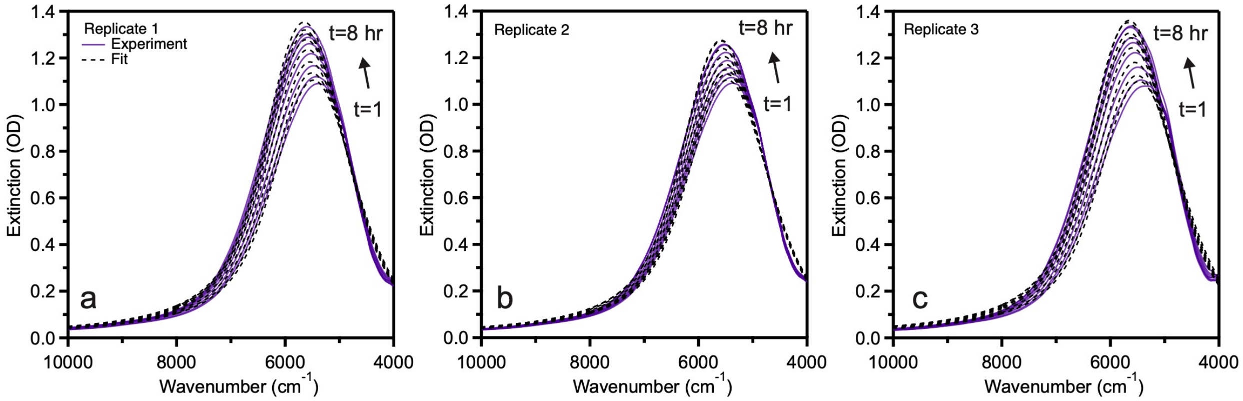


**Figure S10.** Statistical variability and error in EET rates from biological replicates in a single experiment. Three independently grown colonies of *S. oneidensis* MR-1 were dispersed in three cuvettes so spectra could be collected simultaneously. Results from (a) cuvette 1, (b) cuvette 2, and (c) cuvette 3 illustrate that there is variability in optical change among the replicates, but our model is equally able to fit each replicate. We note that cuvette 3 was actually half the pathlength (0.5 mm) of cuvettes 1 and 2, so the extinction was doubled in panel c to more easily compare to panels a and b.

**Table S6. Testing fit robustness against initial guess variability.** Ten sets of values for fit parameters were randomly generated between the fit constraints shown in the table below. All initial guesses converged on the same set of solutions.

| Fit Parameter | $n_e$ | $\sigma_{n_e}$ | $f_e$ |
|---|---|---|---|
| Lower Bound | 1.0E+18 | 1.0E+17 | 0 |
| Upper Bound | 3.0E+21 | 1.0E+21 | 1 |
| Randomly Generated Initial Guess | 9.8E+18 | 1.1E+19 | 0.66 |
| | 1.5E+21 | 1.6E+20 | 0.52 |
| | 7.4E+20 | 3.8E+19 | 0.97 |
| | 1.1E+21 | 2.1E+18 | 0.65 |
| | 1.9E+21 | 2.6E+20 | 0.80 |
| | 1.7E+21 | 1.6E+19 | 0.45 |
| | 1.7E+21 | 8.0E+20 | 0.43 |
| | 8.6E+20 | 5.6E+20 | 0.82 |
| | 1.7E+21 | 3.6E+20 | 0.08 |
| | 3.4E+20 | 7.0E+19 | 0.13 |
| Result | 1.2E+21 | 2.1E+20 | 0.61 |

**Table S7.** HEDA fit results and calculated values quantifying electron transfer extracted from spectra plotted in main text Figure 4. Variables are defined and the standard deviation in each parameter is calculated as described in Note S2.

**Strain:** *S. oneidensis* MR-1

| | Fit Results | | | Calculated Parameters | | |
|---|---|---|---|---|---|---|
| Time (min) | $\mu_{n_e}$ (x $10^{20}$ $cm^3$) | $\sigma_{n_e}$ (x $10^{20}$ $cm^3$) | $F_{v,core}$ | $N_{NC}$ | $\Delta N_{NC}$ | $\Delta N_{\boldsymbol{cell}}$ (x $10^7$) |
| 0 | 10.8 + 0.03 | 2.3 + 0.02 | 0.563 + 0.006 | 492 + 5 | 0 | 0 |
| 60 | 10.8 + 0.06 | 2.2 + 0.02 | 0.616 + 0.012 | 540 + 11 | 49 + 12 | 8.37 + 2.1 |
| 120 | 10.9 + 0.05 | 2.2 + 0.01 | 0.647 + 0.015 | 572 + 13 | 81 + 14 | 13.8 + 2.4 |
| 180 | 11.0 + 0.04 | 2.2 + 0.02 | 0.671 + 0.015 | 595 + 13 | 104 + 14 | 17.8 + 2.5 |
| 240 | 11.0 + 0.03 | 2.2 + 0.01 | 0.684 + 0.011 | 609 + 10 | 117 + 11 | 20.0 + 2.0 |
| 300 | 11.0 + 0.04 | 2.2 + 0.02 | 0.696 + 0.011 | 620 + 10 | 129 + 11 | 22.0 + 2.0 |
| 360 | 11.0 + 0.03 | 2.2 + 0.02 | 0.708 + 0.008 | 630 + 7 | 139 + 9 | 23.8 + 1.6 |
| 420 | 11.0 + 0.04 | 2.2 + 0.02 | 0.718 + 0.008 | 638 + 7 | 147 + 9 | 25.2 + 1.6 |
| 480 | 11.0 + 0.05 | 2.2 + 0.02 | 0.725 + 0.009 | 645 + 9 | 153 + 10 | 26.2 + 1.8 |

**Strain:** *S. oneidensis* S2933

| | Fit Results | | | Calculated Parameters | | |
|---|---|---|---|---|---|---|
| Time (min) | $\mu_{n_e}$ (x $10^{20}$ $cm^3$) | $\sigma_{n_e}$ (x $10^{20}$ $cm^3$) | $F_{v,core}$ | $N_{NC}$ | $\Delta N_{NC}$ | $\Delta N_{\boldsymbol{cell}}$ (x $10^7$) |
| 0 | 10.9 + 0.07 | 2.3 + 0.02 | 0.554 + 0.012 | 487 + 11 | 0 | 0 |
| 60 | 10.9 + 0.06 | 2.2 + 0.02 | 0.591 + 0.018 | 519 + 16 | 33 + 20 | 5.64 + 3.4 |
| 120 | 10.9 + 0.06 | 2.2 + 0.01 | 0.607 + 0.024 | 535 + 22 | 48 + 24 | 8.30 + 4.1 |
| 180 | 10.9 + 0.05 | 2.2 + 0.01 | 0.622 + 0.029 | 549 + 25 | 63 + 28 | 10.7 + 4.7 |
| 240 | 10.9 + 0.05 | 2.2 + 0.01 | 0.637 + 0.030 | 563 + 27 | 76 + 29 | 13.1 + 5.0 |
| 300 | 10.9 + 0.06 | 2.2 + 0.01 | 0.651 + 0.031 | 576 + 28 | 89 + 30 | 15.3 + 5.1 |
| 360 | 10.9 + 0.08 | 2.2 + 0.01 | 0.666 + 0.034 | 588 + 31 | 102 + 33 | 17.4 + 5.6 |
| 420 | 10.9 + 0.10 | 2.2 + 0.01 | 0.676 + 0.034 | 597 + 31 | 110 + 33 | 18.9 + 5.6 |
| 480 | 10.9 + 0.11 | 2.2 + 0.02 | 0.683 + 0.035 | 604 + 32 | 117 + 33 | 2.00 + 5.7 |

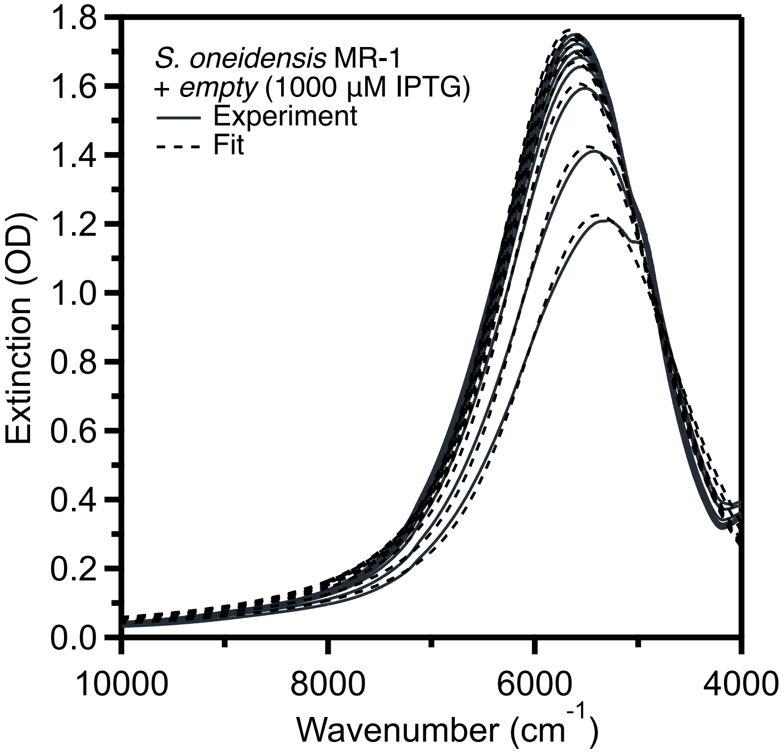


**Figure S11.** *In situ* extinction spectra and HEDA fits for PAAPEO-ITO during EET from *S. oneidensis* MR-1 with an empty vector and 1000 µM inducing molecule (IPTG).

**Table S8.** HEDA fit results and calculated values quantifying electron transfer extracted from spectra plotted in main text Figure 5 and Figure S9. Variables are defined and the standard deviation in each parameter is calculated as described in Note S2.

**Strain:** *S. oneidensis* MR-1 + $empty$ (1000 µM IPTG)

| | Fit Results | | | Calculated Parameters | | |
|---|---|---|---|---|---|---|
| Time (min) | $\mu_{n_e}$ (x $10^{20}$ cm$^3$) | $\sigma_{n_e}$ (x $10^{20}$ cm$^3$) | $F_{v,core}$ | $N_{NC}$ | $\Delta N_{NC}$ | $\Delta N_{\boldsymbol{cell}}$ (x $10^7$) |
| 0 | 10.8 ± 0.07 | 2.2 ± 0.02 | 0.574 ± 0.011 | 501 ± 10 | 0 | 0 |
| 60 | 10.9 ± 0.01 | 2.1 ± 0.01 | 0.625 ± 0.019 | 552 ± 16 | 51 ± 19 | 8.73 ± 3.3 |
| 90 | 11.0 ± 0.03 | 2.0 ± 0.03 | 0.651 ± 0.024 | 579 ± 21 | 78 ± 23 | 13.4 ± 4.0 |
| 120 | 11.1 ± 0.03 | 2.0 ± 0.03 | 0.671 ± 0.022 | 602 ± 20 | 101 ± 23 | 17.3 ± 3.9 |
| 180 | 11.2 ± 0.02 | 1.9 ± 0.02 | 0.692 ± 0.018 | 626 ± 17 | 125 ± 20 | 21.4 ± 3.4 |
| 240 | 11.2 ± 0.02 | 1.9 ± 0.01 | 0.702 ± 0.017 | 637 ± 15 | 136 ± 18 | 23.3 ± 3.1 |
| 300 | 11.2 ± 0.01 | 1.9 ± 0.01 | 0.708 ± 0.016 | 644 ± 14 | 143 ± 18 | 24.5 ± 3.0 |
| 360 | 11.2 ± 0.01 | 1.9 ± 0.01 | 0.716 ± 0.016 | 651 ± 14 | 151 ± 18 | 25.8 ± 3.1 |
| 420 | 11.3 ± 0.01 | 1.9 ± 0.01 | 0.723 ± 0.017 | 658 ± 16 | 157 ± 19 | 26.9 ± 3.2 |
| 480 | 11.3 ± 0.02 | 1.9 ± 0.01 | 0.728 ± 0.019 | 663 ± 18 | 162 ± 20 | 27.8 ± 3.5 |

**Strain:** *S. oneidensis* $\Delta mtrC\Delta omcA\Delta mtrF$ + $mtrC$ (1000 μM IPTG)

| | Fit Results | | | Calculated Parameters | | |
|---|---|---|---|---|---|---|
| Time (min) | $\mu_{n_e}$ (x 10[20] cm3) | $\sigma_{n_e}$ (x 10[20] cm3) | $F_{v,core}$ | $N_{NC}$ | $\Delta N_{NC}$ | $\Delta N_{\boldsymbol{cell}}$ (x $10^7$) |
| 0 | 10.8 ± 0.07 | 2.2 ± 0.04 | 0.571 ± 0.007 | 499 ± 7 | 0 | 0 |
| 60 | 10.8 ± 0.08 | 2.1 ± 0.04 | 0.592 ± 0.012 | 519 ± 11 | 20 ± 13 | 3.43 ± 2.3 |
| 90 | 10.9 ± 0.08 | 2.1 ± 0.04 | 0.595 ± 0.012 | 523 ± 11 | 24 ± 13 | 4.10 ± 2.2 |
| 120 | 10.9 ± 0.07 | 2.1 ± 0.04 | 0.599 ± 0.012 | 527 ± 11 | 28 ± 13 | 4.80 ± 2.2 |
| 180 | 10.9 ± 0.07 | 2.1 ± 0.04 | 0.608 ± 0.013 | 536 ± 12 | 37 ± 14 | 6.28 ± 2.4 |
| 240 | 10.9 ± 0.05 | 2.1 ± 0.04 | 0.616 ± 0.015 | 545 ± 13 | 46 ± 15 | 7.82 ± 2.6 |
| 300 | 10.9 ± 0.05 | 2.1 ± 0.04 | 0.626 ± 0.016 | 555 ± 14 | 55 ± 16 | 9.48 ± 2.7 |
| 360 | 11.0 ± 0.04 | 2.1 ± 0.04 | 0.636 ± 0.017 | 564 ± 15 | 65 ± 17 | 11.1 ± 2.8 |
| 420 | 11.0 ± 0.04 | 2.1 ± 0.04 | 0.646 ± 0.017 | 573 ± 15 | 74 ± 16 | 12.6 ± 2.8 |
| 480 | 11.0 ± 0.04 | 2.1 ± 0.04 | 0.654 ± 0.016 | 580 ± 14 | 81 ± 16 | 13.8 ± 2.7 |

**Strain:** *S. oneidensis* $\Delta mtrC\Delta omcA\Delta mtrF$ + $mtrC$ (100 μM IPTG)

| | Fit Results | | | Calculated Parameters | | |
|---|---|---|---|---|---|---|
| Time (min) | $\mu_{n_e}$ (x 10[20] cm3) | $\sigma_{n_e}$ (x 10[20] cm3) | $F_{v,core}$ | $N_{NC}$ | $\Delta N_{NC}$ | $\Delta N_{\boldsymbol{cell}}$ (x $10^7$) |
| 0 | 10.8 ± 0.04 | 2.2 ± 0.04 | 0.569 ± 0.007 | 498 ± 7 | 0 | 0 |
| 60 | 10.9 ± 0.04 | 2.1 ± 0.03 | 0.585 ± 0.009 | 513 ± 8 | 16 ± 11 | 2.68 ± 1.8 |
| 90 | 10.9 ± 0.03 | 2.1 ± 0.03 | 0.589 ± 0.008 | 517 ± 8 | 20 ± 10 | 3.34 ± 1.7 |
| 120 | 10.9 ± 0.03 | 2.1 ± 0.03 | 0.592 ± 0.007 | 520 ± 7 | 23 ± 9 | 3.91 ± 1.6 |
| 180 | 10.9 ± 0.03 | 2.1 ± 0.03 | 0.597 ± 0.006 | 526 ± 5 | 28 ± 9 | 4.82 ± 1.5 |
| 240 | 10.9 ± 0.03 | 2.1 ± 0.03 | 0.603 ± 0.005 | 532 ± 4 | 35 ± 8 | 5.92 ± 1.4 |
| 300 | 10.9 ± 0.03 | 2.1 ± 0.03 | 0.609 ± 0.004 | 539 ± 4 | 41 ± 8 | 7.04 ± 1.3 |
| 360 | 10.9 ± 0.04 | 2.1 ± 0.03 | 0.618 ± 0.004 | 547 ± 5 | 50 ± 8 | 8.50 ± 1.4 |
| 420 | 11.0 ± 0.04 | 2.0 ± 0.03 | 0.627 ± 0.007 | 555 ± 7 | 58 ± 10 | 9.89 ± 1.6 |
| 480 | 11.0 ± 0.05 | 2.0 ± 0.02 | 0.633 ± 0.008 | 562 ± 8 | 64 ± 10 | 11.0 ± 1.7 |

**Strain:** *S. oneidensis* $\Delta mtrC \Delta omcA \Delta mtrF$ + $mtrC$ (10 μM IPTG)

| | Fit Results | | | Calculated Parameters | | |
|---|---|---|---|---|---|---|
| Time (min) | $\mu_{n_e}$ (x $10^{20}$ $cm^3$) | $\sigma_{n_e}$ (x $10^{20}$ $cm^3$) | $F_{v,core}$ | $N_{NC}$ | $\Delta N_{NC}$ | $\Delta N_{cell}$ (x $10^7$) |
| 0 | 10.7 ± 0.05 | 2.2 ± 0.02 | 0.591 ± 0.008 | 511 ± 8 | 0 | 0 |
| 60 | 10.7 ± 0.03 | 2.2 ± 0.02 | 0.597 ± 0.005 | 518 ± 5 | 7 ± 9 | 1.23 ± 1.5 |
| 90 | 10.7 ± 0.03 | 2.2 ± 0.02 | 0.598 ± 0.005 | 520 ± 5 | 9 ± 9 | 1.53 ± 1.5 |
| 120 | 10.7 ± 0.03 | 2.2 ± 0.02 | 0.599 ± 0.006 | 521 ± 5 | 10 ± 9 | 1.79 ± 1.6 |
| 180 | 10.8 ± 0.02 | 2.2 ± 0.02 | 0.602 ± 0.006 | 524 ± 6 | 13 ± 9 | 2.27 ± 1.6 |
| 240 | 10.8 ± 0.02 | 2.2 ± 0.02 | 0.604 ± 0.006 | 527 ± 5 | 16 ± 9 | 2.80 ± 1.6 |
| 300 | 10.8 ± 0.02 | 2.2 ± 0.03 | 0.606 ± 0.005 | 530 ± 5 | 19 ± 9 | 3.33 ± 1.6 |
| 360 | 10.8 ± 0.02 | 2.2 ± 0.03 | 0.610 ± 0.006 | 533 ± 5 | 23 ± 9 | 3.92 ± 1.6 |
| 420 | 10.8 ± 0.03 | 2.1 ± 0.03 | 0.613 ± 0.006 | 537 ± 6 | 27 ± 9 | 4.56 ± 1.6 |
| 480 | 10.8 ± 0.03 | 2.1 ± 0.04 | 0.617 ± 0.007 | 542 ± 6 | 31 ± 10 | 5.30 ± 1.7 |

**Strain:** *S. oneidensis* $\Delta mtrC \Delta omcA \Delta mtrF$ + $mtrC$ (0 μM IPTG)

| | Fit Results | | | Calculated Parameters | | |
|---|---|---|---|---|---|---|
| Time (min) | $\mu_{n_e}$ (x $10^{20}$ $cm^3$) | $\sigma_{n_e}$ (x $10^{20}$ $cm^3$) | $F_{v,core}$ | $N_{NC}$ | $\Delta N_{NC}$ | $\Delta N_{cell}$ (x $10^7$) |
| 0 | 10.8 ± 0.05 | 2.2 ± 0.01 | 0.577 ± 0.007 | 503 ± 6 | 0 | 0 |
| 60 | 10.8 ± 0.05 | 2.2 ± 0.01 | 0.583 ± 0.006 | 509 ± 6 | 6 ± 9 | 1.04 ± 1.5 |
| 90 | 10.8 ± 0.05 | 2.1 ± 0.01 | 0.583 ± 0.006 | 510 ± 6 | 7 ± 9 | 1.20 ± 1.5 |
| 120 | 10.8 ± 0.04 | 2.1 ± 0.01 | 0.584 ± 0.006 | 511 ± 5 | 8 ± 8 | 1.32 ± 1.5 |
| 180 | 10.8 ± 0.04 | 2.1 ± 0.01 | 0.585 ± 0.006 | 513 ± 5 | 10 ± 8 | 1.66 ± 1.4 |
| 240 | 10.8 ± 0.04 | 2.1 ± 0.01 | 0.586 ± 0.005 | 514 ± 5 | 11 ± 8 | 1.92 ± 1.4 |
| 300 | 10.9 ± 0.04 | 2.1 ± 0.01 | 0.588 ± 0.005 | 516 ± 5 | 13 ± 8 | 2.17 ± 1.4 |
| 360 | 10.9 ± 0.04 | 2.1 ± 0.01 | 0.589 ± 0.005 | 517 ± 5 | 14 ± 8 | 2.38 ± 1.4 |
| 420 | 10.9 ± 0.04 | 2.1 ± 0.01 | 0.590 ± 0.005 | 519 ± 5 | 15 ± 8 | 2.65 ± 1.4 |
| 480 | 10.9 ± 0.04 | 2.1 ± 0.01 | 0.591 ± 0.005 | 520 ± 5 | 17 ± 8 | 2.90 ± 1.4 |

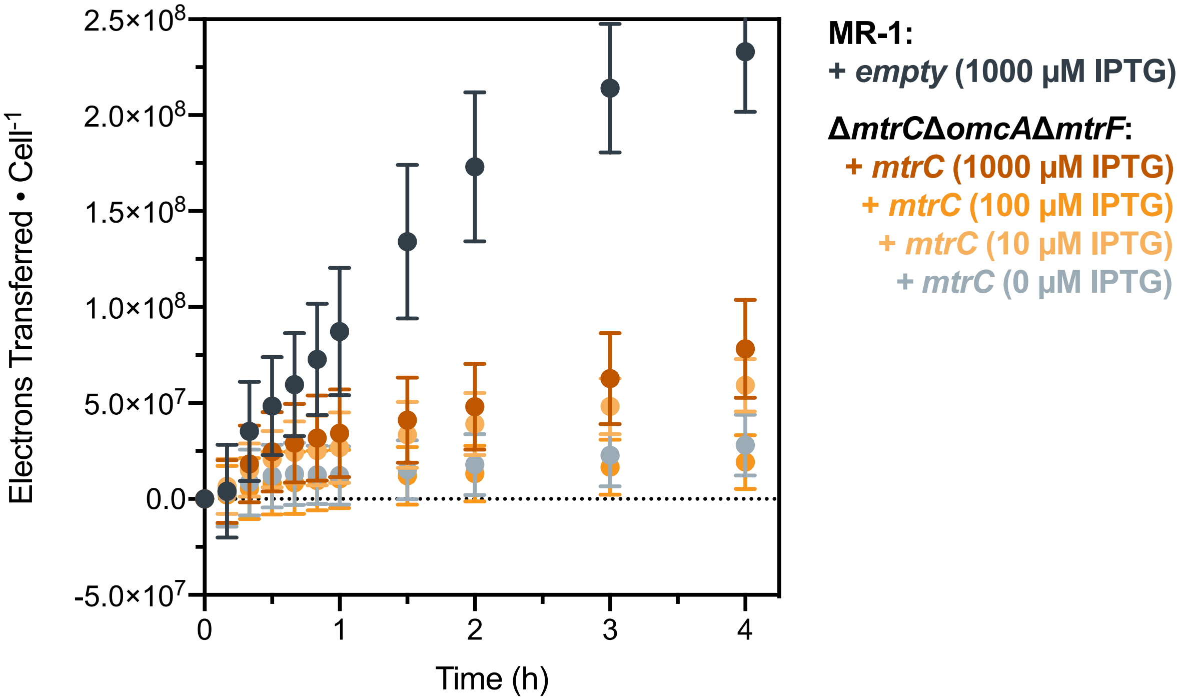


**Figure S12.** Time-resolved spectroscopic measurement of PAAPEO-ITO during EET from *S. oneidensis* Δ*mtrC*Δ*omcA*Δ*mtrF* + *mtrC* and *S. oneidensis* MR-1 + *empty* demonstrates 2 regimes. The early domain (≤ 1 h) shows deviation from linearity due to aerobic respiration, introduction of a deuterated solvent, and cell stress. This regime includes a lag phase (0–20 min) followed by a sharp increase in electron transfer (20–60 min). The early domain is followed by steady-state electron transfer (1–8 h), which was used for fitting analysis of all samples except *S. oneidensis* MR-1 + *empty*, as this sample showed signs of nanocrystal saturation after 3 h. Data shown are mean ± S.D. of $n$ = 3 biological replicates.

**Note S4: Gene expression modeling for EET rate constants.** EET rate constants ($k_{EET}$) were fit to an activating Hill Function model of gene expression, which takes the form:

$$y = Min + (Max - Min)\frac{[I]^n}{{K_{1/2}}^n + [I]^n}, \qquad \text{Equation S7}$$

where *Min* and *Max* refer to the output ($y$) bounds, [$I$] refers to the concentration of inducer (in this system, IPTG), $n$ refers to the hillslope of the fit, and $K_{1/2}$ refers to the concentration of inducer at half-maximal response. The fit parameters obtained from this model ($n$ = 0.9, $K_{1/2}$ = 40 μM) are in agreement with previous studies using this strain, but do not include enough data for reputable fit statistics (*e.g.*, only four data points).[13,14]
